# Mps1 promotes poleward chromosome movements in meiotic pro-metaphase

**DOI:** 10.1101/2020.06.28.176370

**Authors:** Régis E Meyer, Aaron R Tipton, Gary J Gorbsky, Dean S Dawson

**Affiliations:** Program in Cell Cycle and Cancer Biology, Oklahoma Medical Research Foundation, Oklahoma City, OK 73104, USA; Department of Cell Biology, University of Oklahoma Health Sciences Center, Oklahoma City, OK 73104, USA

## Abstract

In prophase of meiosis I, homologous partner chromosomes pair and become connected by crossovers. Chiasmata, the connections formed between the partners enable the chromosome pair, called a bivalent, to attach as a single unit to the spindle. When the meiosis I spindle forms in prometaphase, most bivalents are associated with a single spindle pole and go through a series of oscillations on the spindle, attaching to and detaching from microtubules until the partners of the bivalent are bi-oriented, that is, attached to microtubules from opposite sides of the spindle, and prepared to be segregated at anaphase I. The conserved, kinetochore-associated kinase, Mps1, is essential for the bivalents to be pulled by microtubules across the spindle in prometaphase. Here we show that *MPS1* is not required for kinetochores to attach microtubules but instead is necessary to trigger the migration of microtubule-attached kinetochores towards the poles. Our data support the model that Mps1 triggers depolymerization of microtubule ends once they attach to kinetochores in prometaphase. Thus, Mps1 acts at the kinetochore to co-ordinate the successful attachment of a microtubule and the triggering of microtubule depolymerization to move the chromosome.

## INTRODUCTION

In many organisms, cells enter prometaphase of meiosis with kinetochore-microtubule attachments that would lead to segregation errors if they were not corrected (Meyer et al., 2013; Chmátal et al., 2015; Nicklas, 1997). In budding yeast each partner chromosome in the homolog pair (called a bivalent) can attach one microtubule to its kinetochore (Winey et al., 2005; Sarangapani et al., 2014). The bivalents begin meiosis mono-oriented (both partners at one pole) and, through a series of steps, become bi-oriented and prepared to separate away from each other at anaphase I (Fig. 1 A). The microtubule-organizing centers, called spindle pole bodies in yeast (SPBs), are duplicated in pre-meiotic S-phase resulting in an older SPB and a newly formed SPB. In late prophase the homologous chromosome pairs (called bivalents) cluster at the side-by-side SPBs in a microtubule dependent manner (Fig. 1 A). The end of prophase and entry into pro-metaphase is marked by the formation of a spindle between the SPBs forcing them apart with the bivalents attached mainly to the older SPB. The bivalents are released from this monopolar attachment in an Aurora B-dependent manner (Monje-Casas et al., 2007; Meyer et al., 2013) as was previously demonstrated in mitotic cells (Biggins et al., 1999; Cheeseman et al., 2002; Tanaka et al., 2002). Then, following a series of migrations back and forth across the spindle that include a series of microtubule releases (via Aurora B) and re-attachments the partners of the bivalent become attached to microtubules from opposite SPBs (Meyer et al., 2013). During this process, the spindle assembly checkpoint senses the state of kinetochore-microtubule attachments and delays cell cycle progression into anaphase until all chromosome pairs are bi-oriented (Shonn, 2000; Cheslock et al., 2005).

**Figure 1.**
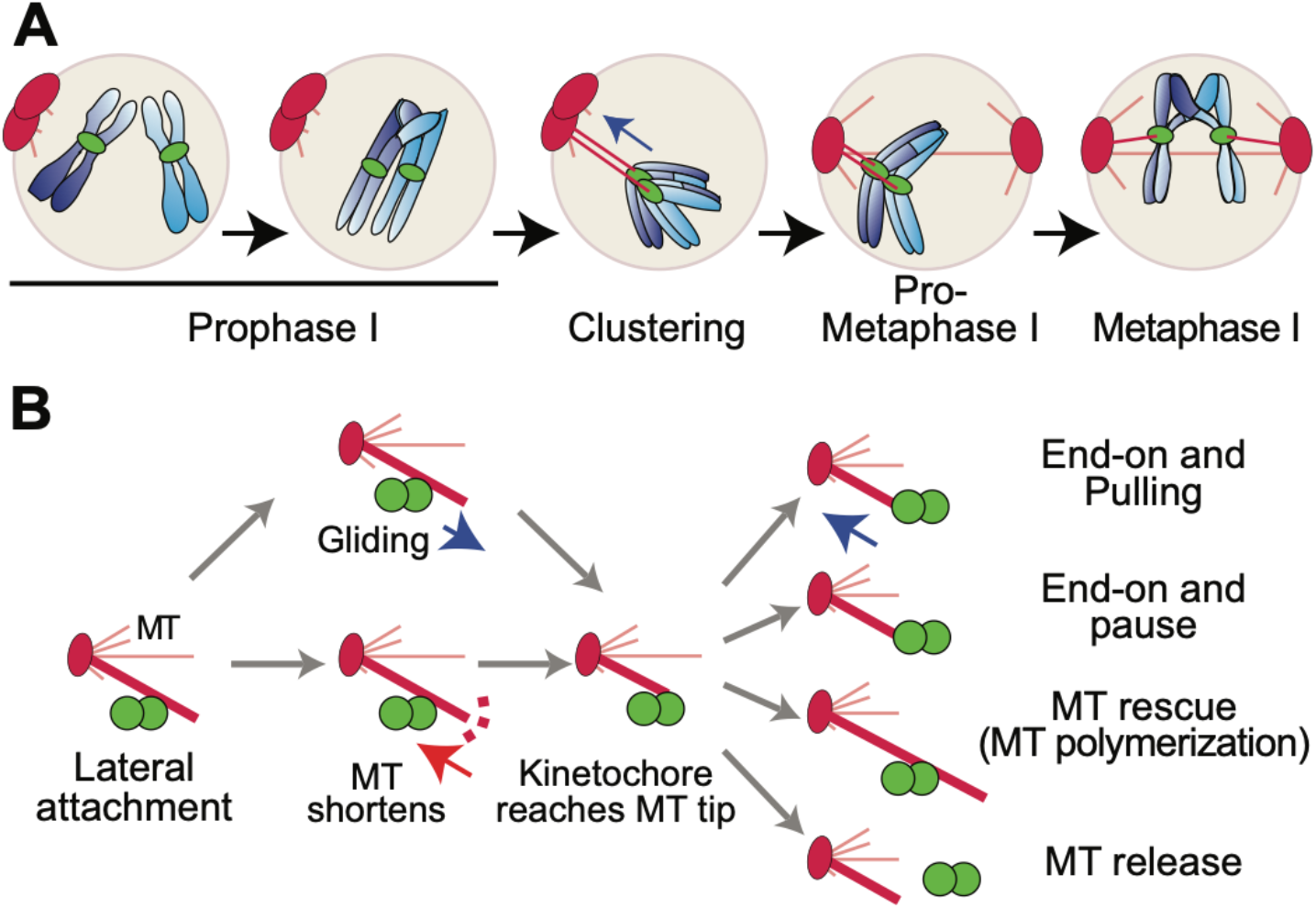
Kinetochore-microtubule interactions in budding yeast meiosis. **A.** In prophase I, chromosomes have released their attachments to microtubules. At the exit from prophase I, centromeres cluster at the side-by-side spindle pole bodies. When spindle pole bodies separate to form a spindle most centromeres are attached to the older spindle pole body. Following a period of oscillations on the spindle including microtubule releases and re-attachments, the homologous partners become bi-oriented. **B.** Studies in mitotic cells suggest most initial attachments are lateral (adapted from (Tanaka, 2010)). Microtubules depolymerize until they meet the kinetochore. In some organisms, kinetochores can glide towards the microtubule plus end. When the microtubule plus end meets the kinetochore, the illustrated outcomes have been observed.

The process of attaching the kinetochores to microtubules appears to be controlled at several levels (reviewed in (Godek et al., 2015; Lampson and Grishchuk, 2017; Tanaka, 2010)). A series of studies from the Tanaka laboratory defined these steps in yeast mitosis (Fig. 1 B). They found that, in yeast, as in other organisms, the kinetochores first attach most often to lateral surfaces of microtubules (Franco et al., 2007; Gachet et al., 2008; Hayden et al., 1990; Magidson et al., 2011; Merdes and De Mey, 1990; Rieder and Alexander, 1990; Tanaka et al., 2005). Second, the microtubule depolymerizes to bring the microtubule plus-end to the kinetochore (Kitamura et al., 2007; Tanaka et al., 2007). The kinetochore and microtubule plus-end can then have any of several fates (Fig. 1 B). The microtubule can re-polymerize, the kinetochore can release the microtubule, or the kinetochore can form an end-on attachment that can move the kinetochore poleward as the microtubule de-polymerizes. In this process, the protein composition at the kinetochore-microtubule interface, and modifications of those proteins, change, which promotes the ability of the kinetochore to track the shortening microtubule (Asbury et al., 2006; Daum et al., 2009; Gaitanos et al., 2009; Grishchuk et al., 2008; Lampert et al., 2010; Powers et al., 2009; Schmidt et al., 2012; Umbreit et al., 2014; Volkov et al., 2013; Welburn et al., 2009; Westermann et al., 2006).

Mps1 is a conserved kinase with a central role in the spindle assembly checkpoint (Weiss and Winey, 1996; Hardwick et al., 1996; Abrieu et al., 2001). In budding yeast, Mps1 also has an essential role in meiotic chromosome segregation (Straight et al., 2000) where it is necessary for the efficient formation of force-generating attachments of kinetochores to microtubules that result in processive poleward migration during the bi-orientation process (Fig. 1 A) (Meyer et al., 2013). In *MPS1* mutants, following anaphase I, most chromosomes end up associated with the spindle pole with which they were initially associated when the spindle first formed (Meyer et al., 2013). This is because they cannot move across the spindle to the opposite pole in pro-metaphase. Since most chromosomes connect to the older SPB just before pro-metaphase, even in wild-type cells, *MPS1* mutants exhibit over 80% non-disjunction, nearly all to the older SPB at anaphase I.

This role of Mps1 in promoting force-generating kinetochore-microtubule attachments is critical for meiosis but less so in mitosis (Meyer et al., 2013). In budding yeast as in many other organisms, *MPS1* is an essential gene, but separation-of-function alleles have been identified that result in severe defects in meiotic bi-orientation but very mild defects in mitosis (Meyer et al., 2013). This suggests either that meiosis is particularly sensitive to defects in the bi-orientation machinery, or alternatively, meiotic sensitivity to *MPS1* mutations reflects a meiosis-specific process. Interestingly, similar meiosis-specific mutant alleles of *MPS1* have also been isolated *Drosophila* and zebrafish (Gilliland et al., 2005; Poss et al., 2004).

The manner in which Mps1 promotes the formation of force-generating attachments between kinetochores and microtubule plus-ends is unclear. Does Mps1 promote the movement of kinetochores towards the spindle mid-zone so they can encounter microtubules from the opposite pole, or convert lateral attachments to end-on attachments, or stabilize end-on kinetochore-microtubule attachments, or depolymerize microtubules to drag kinetochores poleward (Fig. 1 B)? Because Mps1 kinase is known to have many targets, it could be involved in coordinating multiple steps in the bi-orientation process. Here we use live cell imaging experiments to explore the meiotic roles of Mps1. The results of these experiments suggest that *MPS1* mutants can form end-on kinetochore-microtubule attachments but demonstrate that they are defective in the subsequent microtubule depolymerization that pulls kinetochores poleward.

## MATERIALS AND METHODS

### Yeast strains and culture conditions

All strains are derivatives of two strains termed X and Y described previously (Dresser et al., 1994). We used standard yeast culture methods (Amberg et al., 2005). To induce meiosis, cells were grown in YP acetate to 4-4.5×10^7^ cells per ml, and then shifted to 1% potassium acetate at 10^8^ cells per ml. Mitotic cells were grown in SD-TRP media (Sunrise Science).

### Genome modifications

Heterozygous and homozygous *CEN1-GFP* dots: An array of 256 *lac* operon operator sites on plasmid pJN2 was integrated near the *CEN1* locus (coordinates 153583–154854). *lacI-GFP* fusions under the control of *P*_*CYC1*_ and *P*_*DMC1*_ were also expressed in this strain to visualize the location of the *lacO* operator sites during meiosis as described in (Meyer et al., 2013)).

PCR-based methods were used to create complete deletions of ORFs and promoter insertions (Janke et al., 2004; Longtine et al., 1998). *spo11::KANMX, spo11::HIS3MX6, P_GPD1_-GAL4(848)-ER-URA3::hphNT1, natNT2::P_GAL1_-NDT80*, *KANMX::P_GAL1_-NDT80, mps1::KANMX, TRP1::10Xmyc-mps1-as1 (=mps1-as1), mps1-R170S::his5, KANMX::P_CLB2_-3HA-MPS1 (=mps1-md)*, *KANMX::P_CLB2_-3HA-NDC80 (=ndc80-md), KANMX::P_CLB2_-3HA-DAM1 (=dam1-md), SPC42-DsRed-URA3* strains were generated previously (Meyer et al., 2018; 2013). The *mEos2-TUB1* strains were generated by inserting *pHIS3p:mEos2-Tub1+3’UTR::TRP1* plasmid (https://www.addgene.org/50652/) in the *TUB1* locus as described (Markus et al., 2015). The *SPC42-GFP-TRP1* strain was a gift from Mike Dresser (as described in (Adams and Kilmartin, 1999)).

### Fluorescence microscopy

Long term live cell imaging experiments (every 45-120 seconds for 3-4 hours) were performed with CellAsic microfluidic flow chambers (www.emdmillipore.com) using Y04D plates with a flow rate of 5 psi. Images were collected with a Nikon Eclipse TE2000-E equipped with the Perfect Focus system, a Roper CoolSNAP HQ2 camera automated stage, an X-cite series 120 illuminator (EXFO) and NIS software. Images were processed and analyzed using NIS software. For the time-lapse imaging of *CEN1* movement, two different exposure programs were defined depending of the presence (*SPO11*) or absence (*spo11Δ*) of chiasmata. In the presence of chiasmata, the intervals were every two minutes for two hours and later every five minutes for two hours (Fig. S1). Without chiasmata, images were acquired every 45 seconds for 75 minutes followed by every 10 minutes for three hours (Fig. 2 and 3).

**Figure 2.**
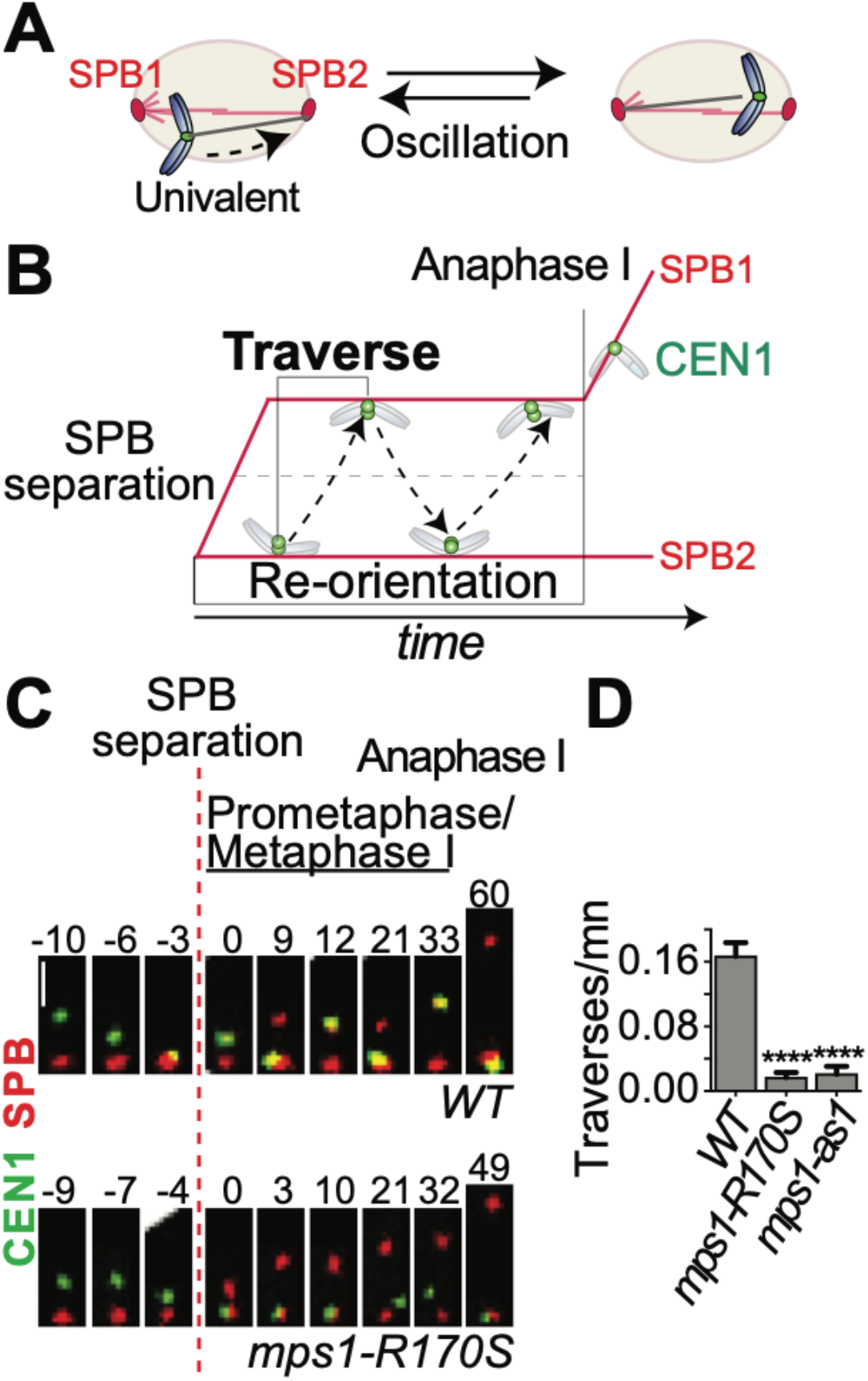
Mps1 promotes processive migration across the meiotic spindle. **A.** Cartoon illustrating the process of re-orientation in the absence of links between homologs (*spo11Δ* background). As the univalent doesn’t have the ability to bi-orient, it will re-orient indefinitely. **B.** The re-orientation process in the *spo11* background can be evaluated by quantifying the traverses across the spindle. **C.** *spo11Δ* diploid cells, with the indicated genotypes, with one GFP-tagged *CEN1* and the SPB marker (*SPC42-DsRed*) were sporulated and released from a pachytene arrest (*P_GAL1_-NDT80 GAL4-ER*) at six hours after meiotic induction by the addition of 5 μM β-estradiol. The cells were observed by a time-lapse movie at 45 second intervals for 75 minutes. Representative kymographs from wild-type and *msp1-R170S* cells are shown. Scale bar: 2 μm. **D.** The number of CEN1-GFP traverses per minute during the first twenty minutes following spindle formation was determined in individual cells. ****p < 0.0001 (Student’s t test). (n≥18).

**Figure 3.**
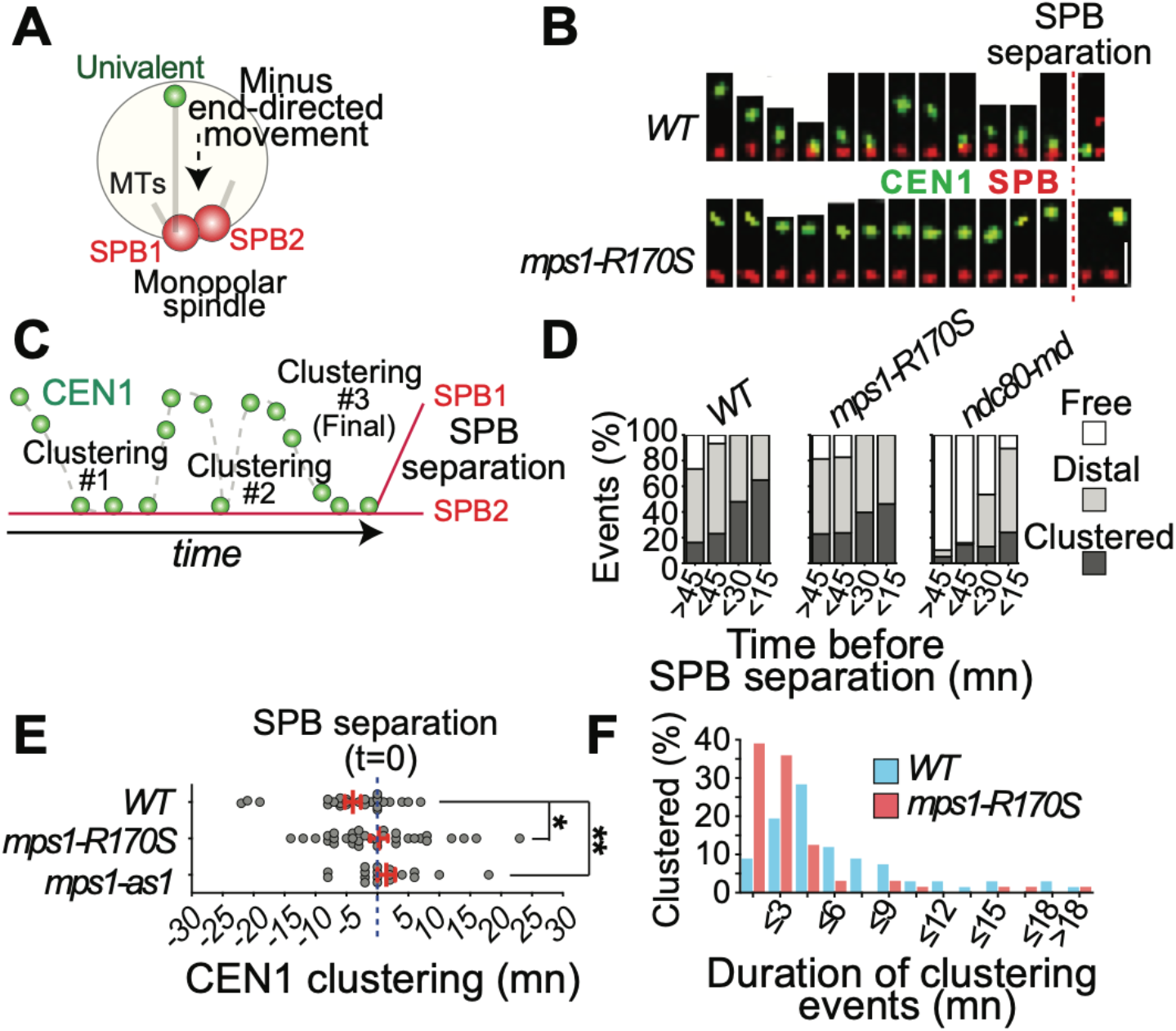
Mps1 promotes minus-end directed migration to the base of monopolar microtubule arrays. **A.** Schematic representation of centromere clustering on a monopolar microtubule array. **B.** *spo11Δ* diploid cells were imaged at 45 second intervals for 75 minutes. Representative wild-type and *mps1-R170S* cells showing clustering, or a failure to cluster, before spindle formation. The dotted line shows the last frame before the spindle formed. Scale bar: 2 μm. **C.** The pulling of the chromosome can be separated in two alternating phases where *CEN1* is either moving towards the SPBs (Clustering) or at a relative constant distance from the SPBs. **D.** The status of *CEN1* was monitored, at each interval, for each individual cell, of the indicated genotypes and classified as follows; Free (white) when the relative position of *CEN1* to the SPBs was un-stable (moving closer and farther from the SPBs in consecutive frames), distal (grey) when *CEN1* was more than 0.5 μm from the SPBs and staying at a constant distance or moving incrementally closer or farther from the SPBs in consecutive frames, and clustered (dark grey), when *CEN1* was staying close (less than 0.5 μm from the SPBs). We assume that most “distal” and “clustered” *CEN1*s are attached to microtubules and “free” *CEN1*s are not. The proportion for each class in each period of 15 minutes preceding bipolar spindle formation is shown (n≥10 cells). **E.** The timing of the final clustering of *CEN1* was monitored relative to the time of SPB separation for each individual cell (n≥19 per genotype). The red dotted line represents the time at which the SPBs separated. *p < 0.05, **p < 0.01 (Student’s t test). **F.** The duration of each clustering event was monitored (number of minutes *CEN1* stayed within 0.5 μm of the SPBs). The distribution of those events in the wild-type and *mps1-R170S* mutant cells (67 vs. 64 events respectively) is shown. The average time spent clustered in *mps1-R170S* cells was significantly less than in wild-type cells (unpaired t test, p = 0.0022).

For monitoring movements of *CEN1-GFP* on monopolar spindles (side-by-side SPBs), following the release from prophase, centromeres were considered as un-attached if they did not remain at a constant distance from the SPBs for at least four consecutives frames. Centromeres were considered to be attached if they stay at a constant distance from the SPBs for at least three consecutive frames or moved incrementally in one direction. The beginning of clustering was defined *CEN1-GFP* first reaches a position within 0.75 μm of the SPB in three consecutive frames. Later events of clustering were defined when centromeres reached, after being attached, a similar position for at least one frame. Traverses (*CEN1* crossing the spindle from one pole to the other one) were counted only when *CEN1-GFP* signal was overlapping with the SPB signal for at least one frame. Homologs were considered to bi-oriented when the the homologous CEN1-GFP signals were distinctly separated in two foci.

For high-speed live cell imaging, images were collected every two seconds for five minutes using a Roper CoolSNAP HQ2 camera on a Zeiss Axio Imager 7.1 microscope fitted with a 100×, NA1.4 plan-Apo objective (Carl Zeiss MicroImaging), an X-cite series 120 illuminator (EXFO), and a BNC555 pulse generator (Berkeley Nucleonics) to synchronize camera exposure with focusing movements and illumination. Cells from sporulating cultures were concentrated, spread across polyethyleneimine-treated coverslips, then covered with a thin 1% agarose pad to anchor the cells to the coverslip. The coverslip was then inverted over a silicone rubber gasket attached to a glass slide. Thru-focus images were acquired as described previously then de-convolved to provide a two-dimensional projected image for each acquisition (Conrad et al., 2008). For the analysis of centromere movements on bipolar spindles, the coordinates of the two SPBs (labeled by *SPC42-GFP*) and the centromeres (marked by *CEN1-GFP*) were defined for each interval. To separate the movement inherent to spindle rotation inside the cells and the movement of *CEN1* on the spindle, a relative position for *CEN1* and the two SPBs was assigned for each interval. For one SPB (SPB1) this position was defined as being constant as x=0 and y=0. For the other SPB (SPB2), the position was defined as x=distance between the SPBs in each frame and y=0. Finally, the relative position of *CEN1* was determined by the distance between *CEN1* and SPB1 and the angle formed between the axis SPB1-SPB2 and SPB1-CEN1. As the acquisitions were done in two dimensions, the impact of the spindle rotating in three dimensions was corrected by assuming the spindle length remains the same or increases over time. Therefore, for instances in which the SPB1-SPB2 distances decreased in sequential frames the value was corrected by replacing the SPB1-SPB2 distance with the prior maximum spindle length (dMax SPB1-SPB2). The magnitude of this correction was also then applied to correct the SPB1-CEN1 distance, the following formula was applied for each interval: Distance SPB1-CEN1 = Observed distance SPB1-CEN1 x dMax SPB1-SPB2 / observed distance SPB1-SPB2. The velocity of *CEN1* movement on the spindle was calculated for each interval by adding the distance between interval n-1 to n+1 and dividing by time interval (4 seconds). The median position for *CEN1* was determined in five-minute intervals for each cell by calculating the average position. The dispersion distance was determined for each interval by calculating the distance between *CEN1* and this average position. Cells with the following characteristics were selected to monitor poleward migration (Fig. 5): The *CEN1* exhibited a migration of 0.9 μm to 1.2 μm to its final destination within 0.25 μm of one SPB. The angle of approach had to be within 15°C on the pole to pole spindle axis. The migrations started within the same half-spindle of the destination SPB. Inside this 0.9-1.2μm-distance movement, the intermediate steps were considered poleward movement when the distance between SPB and *CEN1* from one interval to the other one was decreasing and anti-poleward movement when increasing. The pauses and reversals of direction were determined as follows. First, the distance (D) between the final SPB destination and *CEN1* was calculated for each interval (frame). Second, the average distance for each sequential pair of steps was determined. Third, sequential positions in this sliding average were compared. If the distance between the SPB and *CEN1* was increasing (D≥0), the movement was considered to be paused/reversed. The number of consecutive poleward steps was determined as number of consecutive steps showing continued decreasing distance (D<0).

### Measuring microtubule turnover

Microtubule turnover was evaluated in yeast cells expressing mEos2-Tub1, harvested from either log-phase vegetative cultures (in YPAD medium (Amberg et al., 2005)) or meiotic cultures. For meiotic experiments, cells in a pachytene arrest were induced to exit prophase by the addition of estradiol to the medium, using previously published methods (Meyer et al., 2013). Where indicated, auxin (2 mM, Sigma Aldrich I5148-10G) CuSO4 (200 μM, Sigma Aldrich 451657-10G) or 1-NMPP1 (5 μM, Calbiochem; 5 mM stock in dimethylsulfoxide) were added to the medium at the time of prophase exit. One hour after inducing prophase exit, cells were concentrated, spread across polyethyleneimine-treated coverslips, then covered with a thin 1% agarose pad to anchor the cells to the coverslip. The coverslip was then inverted over a silicone rubber gasket attached to a glass slide. Cells synchronously entering prometaphase were then subjected to imaging to measure microtubule turnover.

Cells were imaged using a 100x, NA 1.4 objective on a Zeiss Axio Observer inverted microscope equipped with a Yokogawa CSU-22 (Yokogawa) spinning disk, Mosaic (digital mirror device, Photonic Instruments/Andor), a Hamamatsu ORCA-Flash4.0LT (Hamamatsu Photonics), and Slidebook software (Intelligent Imaging Innovations). Photoconversion was achieved by targeting a selected area in half the spindle with filtered light from the HBO 100 via the Mosaic, and confocal GFP and RFP images were acquired at 15 sec intervals for ~5 min. At each acquisition, we acquired seven images in the Z-dimension with 0.5 μm spacing. To quantify fluorescence dissipation after photoconversion, we measured pixel intensities within an area surrounding the region of highest fluorescence intensity and background subtracted using an area from the non-converted half spindle using MetaMorph software. Fluorescence values were normalized to the first time-point after photoconversion for each cell and the average intensity at each time point was fit to a single exponential decay curve F = A × exp(-k x t), using SigmaPlot (SYSTAT Software), where A represents the microtubule population with a decay rate of k, respectively. t is the time after photoconversion. For each experiment we performed at least three biological replicates with at least three cells imaged per experiment. Cell numbers for each experiment are in Table 1. Sample identity for scoring fluorescent signals was blinded. The half-life for the microtubule population was calculated as ln2/k. Graphs were prepared using GraphPad Prism. Graphs represent the averages and standard error of the mean for combined replicates.

**Table 1.** Microtubule turnover measurements.

| Diploid strain | Stage | n | Half-time (t1/2; s) | Spindle length (μm) | R2 |
| --- | --- | --- | --- | --- | --- |
| *WT* | Mitotic metaphase | 10 | 119.5 | nd | 0.938 |
| *WT* | Meiotic metaphase | 25 | 119.5 | 2.69 | 0.961 |
| *WT* (1-NMPP1) | Meiotic metaphase | 26 | 126 | 2.62 | 0.986 |
| *mps1-as1* (1-NMPP1) | Meiotic metaphase | 29 | 133.3 | 3.55 | 0.986 |
| *STU2-AID* * | Meiotic metaphase | 29 | 135.9 | 2.46 | 0.973 |
| *STU2-AID* * (+Auxin) | Meiotic metaphase | 23 | 223.6 | 2.15 | 0.945 |
| *spo11* | Meiotic prometaphase | 35 | 79.7 | 3.34 | 0.979 |
| *spo11 mps1-R170S* | Meiotic prometaphase | 25 | 115.5 | 3.11 | 0.932 |
| *spo11* (1-NMPP1) | Meiotic prometaphase | 20 | 80.6 | 3.59 | 0.981 |
| *spo11 mps1-as1* (1-NMPP1) | Meiotic prometaphase | 29 | 119.5 | 3.09 | 0.995 |

## RESULTS

### Mps1 is necessary for chromosome movements across the meiotic spindle

Previous work has shown that Mps1 is necessary for establishing force-generating attachments of kinetochores to microtubules. This is a multi-step process (Fig. 1 B), and the step, or steps, at which *MPS1* mutants are defective is unknown. Therefore, we used live-cell imaging to track chromosome movements at various stages of the meiotic bi-orientation process in order to identify the deficiencies that occur when *MPS1* is inactive.

We focused on the *mps1-R170S* mutation because this separation-of-function allele has only mild mitotic defects and severe meiotic defects, thus providing clues as to the critical roles that Mps1 plays in meiosis. As a control, we used an analog-sensitive allele that allowed us to inactivate the Mps1 kinase activity with an ATP analogue (*mps1-as1*) (Jones et al., 2005). Prior studies revealed that both mutations result in high levels of meiosis I non-disjunction (Meyer et al., 2013). To track chromosome movement, one chromosome (chromosome *I*) was tagged adjacent to its centromere with an array of *lac* operator repeats and the cells expressed lacI-GFP, which binds to the repeats, from a meiotic promotor (Straight et al., 1996). The movement of this GFP-tagged centromere was tracked in cells with a deletion of *SPO11.* In this background, homologous partner chromosomes do not become connected by recombination events to form bivalents (Fig. 2 A) (Klapholz et al., 1985; Loidl et al., 1994). The resulting partnerless univalents, each with only one kinetochore, can never bi-orient on the spindle and thus go through repeated cycles of microtubule attachment, migration on the spindle, and microtubule detachment (Fig. 2 B) (Meyer et al., 2013). Using this assay, both *mps1-as1* and *mps1-R170S* mutants exhibit a nearly complete loss in the ability of chromosomes to traverse across the spindle, while in wild-type cells the GFP-tagged chromosome crosses the spindle, on average, about once every six minutes during pro-metaphase (Fig. 2 C and D).

The coupling of kinetochores to the plus ends of de-polymerizing microtubules is presumably the major driving force for the poleward movements that occur on bipolar spindles. However, in assays with bi-polar spindles (as in Fig. 2 C) it is difficult to know exactly how the kinetochore of a particular chromosome is attached to a microtubule.

The rapid and processive migrations across the mid-zone and to the opposite pole are most consistent with the kinetochore being dragged by a depolymerizing plus-end attached microtubule towards the spindle pole where its plus end is attached (Tanaka et al., 2007). However, it is formally possible that these movements could be gliding of the centromere along the side of a microtubule in the opposite direction, away from the SPB and towards the plus-end of the microtubule it is tracking (Fig. 1 B) (Kapoor et al., 2006; Windecker et al., 2009; Akera et al., 2015).

To distinguish between these possibilities, we assayed the clustering of a univalent chromosome (*spo11* background) towards the side-by-side SPBs before the bipolar spindle is formed (Fig. 1 A and 3 A). At this stage the microtubules form a monopolar array from the side-by-side SPBs, so all poleward movements of chromosomes are minus-end directed and all movements away from the pole are toward the microtubule plus ends. In this experiment, cells were released from a prophase arrest and chromosome movements on the monopolar array were monitored (Fig. 3 A-C). In wild-type cells the univalent migrated towards the side-by-side SPBs (clustering) in consecutive cycles (Fig. 3 B and C) and as cells approached the time of spindle assembly, GFP-tagged centromeres were more and more likely to have become positioned against the SPBs (Fig. 3 D). The beginning of clustering, about thirty minutes before spindle assembly may correspond to the time at which new Ndc80 complexes, capable of interacting with microtubules, are added to the meiotic kinetochore (Meyer et al., 2015; Miller et al., 2012; Chen et al., 2020). This clustering does not occur in *ndc80-md* mutants that cannot produce new outer kinetochores after exiting prophase (Fig. 3 D). The majority of wild-type cells cluster the GFP-tagged centromere five minutes before spindle assembly, while this is significantly delayed in the *MPS1* mutants (Fig. 3 E), and the length of time centromeres remain at the SPB is shorter in *MPS1* mutants (Fig. 3 F). Similar observations were obtained by monitoring bivalent pairs (*SPO11*) (Fig. S1). The data from the clustering experiments support the conclusion that Mps1 is needed to promote movements towards the minus ends of microtubules.

### *mps1-R170S* mutants exhibit pausing defects during the bi-orientation process

The imaging experiments above (and a prior characterization of Mps1 in meiosis, (Meyer et al., 2013)) employ long frame intervals (every 2 minutes) to allow acquisition of data for cells proceeding from pro-metaphase thru anaphase I without photo-bleaching or toxicity. At this frame rate, a traverse of a centromere across the entire spindle can occur in the interval between sequential frames and details about pauses, re-starts and reversals of direction that occur as the kinetochore interacts with a microtubule are not detected. Understanding these details might clarify at which steps in the bi-orientation process Mps1 is playing a critical function. To identify chromosome movements that occur within a single traverse we imaged chromosome behavior at much faster acquisition rates (two second intervals) over the course of five minutes. Images were acquired using a thru-focus method in which a single image is collected as the objective lens focuses thru the cell (Conrad et al., 2008). De-convolution of the acquired data then produces a two-dimensional projection of the image. To reduce acquisition times, the spindle pole bodies (SPBs) and the centromere of chromosome *I* were both tagged with GFP.

Chromosome behavior was quantified in cells with bipolar spindles. In wild-type control cells, chromosomes exhibited several behaviors during the five minute “snapshots” of pro-metaphase. We assigned these behaviors to five categories (Fig. 4 A). These included: i) clustering at one SPB, ii) maintaining a position between the poles (non-polar), iii) low-mobility half spindle -small movements within one half-spindle, iv) high mobility half spindle - directed movements, towards or away from the SPB, in one-half of the spindle without crossing the spindle mid-zone, and v) traverses across the spindle. In most wild-type cells the centromere exhibited at least one traverse or half-spindle length migration in a five-minute window of pro-metaphase (Fig. 4 A, iv and v). These high mobility movements were greatly reduced in *mps1-R170S* mutants (Fig. 4 A, iv and v). In contrast, it was uncommon in the wild-type control strain for centromeres to linger in a non-polar position (Fig. 4 A, ii), but this occurred significantly more frequently in *mps1-R170S* mutants where it was the predominant category.

**Figure 4.**
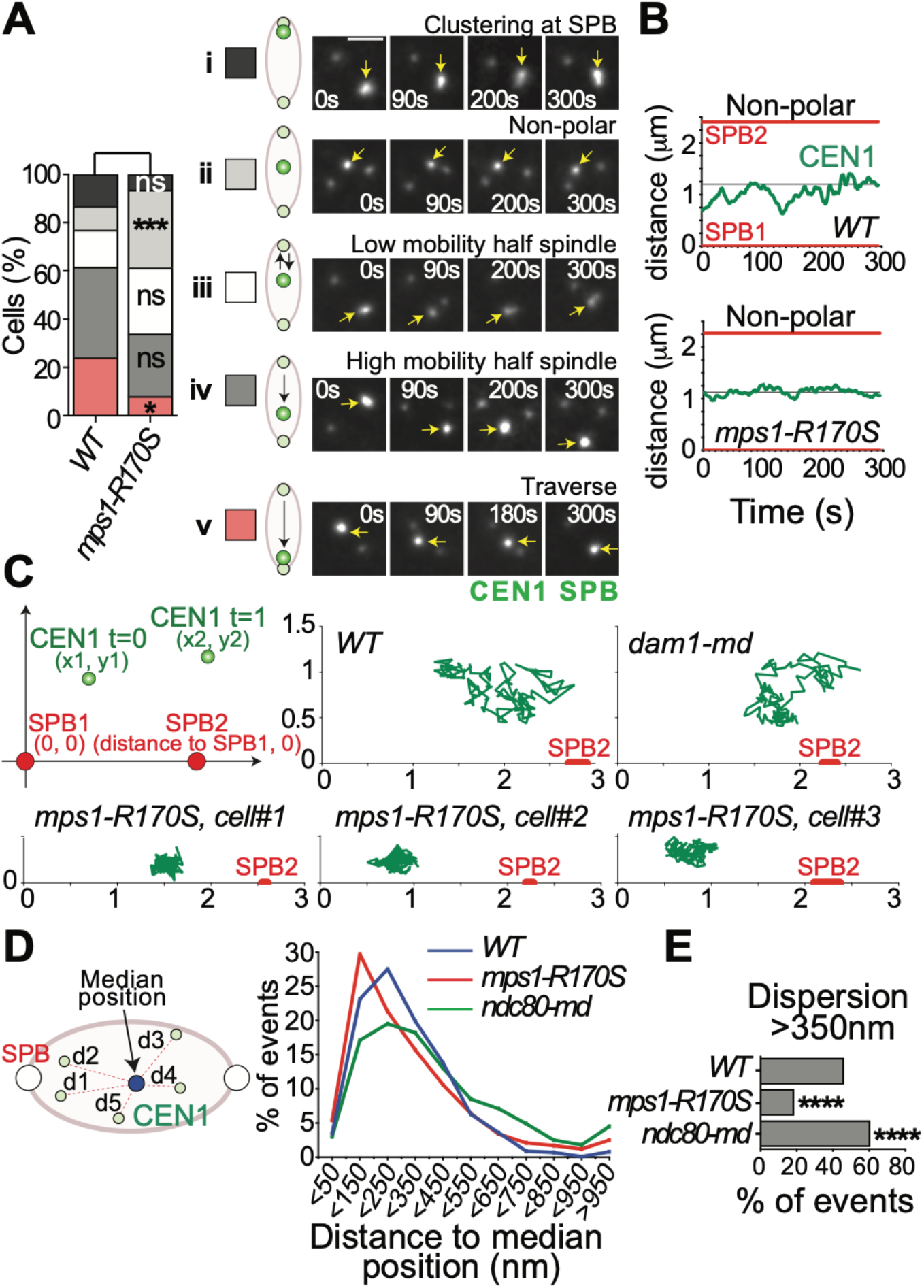
Mps1 promotes chromosome mobility on the meiotic pro-metaphase spindle. *spo11Δ* diploid cells, with the indicated genotypes, with one *CEN1-GFP* tagged chromosome and a SPB marker (*SPC42-GFP*) were sporulated and released from a pachytene arrest (*P_GAL1-_NDT80 GAL4-ER*) at six hours by the addition of 5 μM β- estradiol. Cells were observed by time-lapse imaging during meiosis at two second intervals for five minutes. **A.** According to the primary behavior of the GFP-tagged centromere during the five minute snapshot, cells were placed in one of the following five categories: clustered (remaining close to one SPB), non-polar (positioned away from the poles and not migrating towards a pole), low mobility half spindle (making small movements within on half spindle), high mobility half spindle (moving poleward or toward the mid-one across a half-spindle, traverse (moving pole-to-pole across the spindle). Examples of each classification are shown. Scale bar: 2 μm. **B.** Representative kymographs of wild-type and *mps1-R170S* cells that were classified as “non-polar”. **C.** The top left panel is a schematic of the relative positions of the GFP-tagged *CEN1* in two sequential imaging frames (SPBs are shown in red). The spindle-centered reference system has three key parameters: The position of SPB1 is constant at x=0 and y=0, the position of SPB2 depends on the spindle length (variable over-time) and the coordinates x and y (in microns) define the distance of *CEN1* from SPB1 at that imaging frame. Shown are traces of the location of *CEN1*, relative to the SPBs, in 150 sequential time points (every 2 seconds for 5 minutes) in five representative cells from the non-polar category. **D.** For centromeres classified as non-polar, we calculated the median position of *CEN1* over the course of the five minutes imaging period, and then determined distance of *CEN1* from that median position for each frame of the acquisition. The graph shows the distribution of distances between the centromere and its average position for over 750 frames for each genotype. **E.** The proportion of individual *CEN1* positions more than 350 nm distant from the median position was calculated for each indicated genotype. Mutant genotypes were compared to the wild-type control. ****p < 0.0001 (Fisher’s exact test). n≥750.

Furthermore, the centromeres scored as “non-polar” in *mps1-R170S* cells appeared more stationary than those in wild-type cells (Fig. 4 B). To quantify this, we plotted the positions of the GFP-tagged centromere relative to the SPBs in every frame of the five-minute movie (150 frames) (Fig. 4 C). Representative traces of the GFP-tagged centromeres in a wild-type cell, a *dam1-md* mutant (which is defective in maintaining end-on kinetochore attachments; (Meyer et al., 2018)) and three *mps1-R170S* cells show that in the *mps1-R170S* mutants the centromeres appear locked-in-place (Fig. 4C). We quantified all of the movements of centromeres in the non-polar category (Fig. 4 A ii) by determining the median position of each centromere over the five-minute movie, then determining the distance of the centromere from that position in each of the 150 frames (Fig. 4 D cartoon). The data for wild-type cells, *mps1-R170S* mutant cells and *ndc80-md* mutant cells (in which kinetochores cannot connect to microtubules) are shown in Figure 4 D. This analysis reveals that in *mps1-R170S* cells the centromere stays within a smaller area during prometaphase than is observed in wild-type cells (Fig. 4 D). Furthermore, *mps1-R170S* centromeres exhibit significantly fewer long movements (over 350 nm) – note that the spindle length in these experiments is about 2 μm (Fig. 4 E). *ndc80-md* mutants have the opposite phenotype from *mps1-R170S* mutants – *ndc80-md* centromeres exhibit more movement during prometaphase than is seen in wild-type cells (Fig. 4 D and E). Thus, kinetochore-microtubule attachments restrain the centromeres in prometaphase, perhaps countering the dramatic telomere-led chromosome movements that begin in meiotic prophase and diminish as cells approach metaphase (Conrad et al., 2008; Koszul et al., 2008).

### *mps1-R170S* mutants exhibit reduced processivity during poleward centromere migrations

The static behavior of the non-polar centromeres in *mps1-R170S* mutants is consistent with the model that these represent kinetochores that are attached to the ends of microtubules that are not de-polymerizing. This could be analogous to the “paused” kinetochore-microtubule attachments observed in mitotic budding yeast cells by the Tanaka laboratory (Tanaka et al., 2005; Tanaka, 2010; Tanaka et al., 2007), that sometimes occur when a microtubule depolymerizes until it meets a laterally attached kinetochore (Fig. 1 B). The elevated numbers of the static non-polar centromeres in *mps1-R170S* cells is consistent with the model that one role of Mps1 is to phosphorylate targets at the end-on attached kinetochore-microtubule interface to help converted paused kinetochores to moving kinetochores. To investigate this model, we characterized the behavior of centromeres making poleward migrations in wild-type and *mps1-R170S* cells. We identified centromeres that in the course of our five-minute snapshot of prometaphase moved from a position that was about one micron (0.9 – 1.2 μm) away from a spindle pole towards that pole (Fig. 5 A). Such cells are rare in the *mps1-R170S* population due to the preponderance of locked-in-place centromeres. These poleward migrations could come from either pushing or pulling forces, but since the migrations occur within a half spindle (the average spindle length was over two microns) they are presumably mediated most often by minus-end directed movements along a microtubule that emanates from the destination pole (Fig. 5 A). The chart of the movements of each tracked centromere as it moves poleward (Fig. 5 B) reveals first, that all centromeres exhibit some reversals and pauses during the journey. Some of these might be artifactual as, 1) the measurements are taken from two-dimensional projections of three-dimensional spindles so spindle rotations in the Z-dimension could distort the true kinetochore-SPB distance, and 2) the movements are relatively small compared to the sizes of the centromere GFP and SPB foci – distances measured are from the center of each focus. Measuring protocols were used to minimize these issues (see Methods). Tracking the individual centromeres showed that poleward migrations took significantly less time in wild-type cells than in *mps1-R170S* mutants (Fig. 5 B and C). To determine whether this was because centromeres reach higher velocities in wild-type cells, we measured the velocities of both poleward and anti-poleward centromere movements over the course of migrations to the pole (Fig. 5 D). Measurements were obtained as a sliding three-frame window (four seconds) in which the centromere moved in the same direction between frames one and two and between frames two and three. There was no obvious difference in the average speeds of either poleward or anti-poleward movements of the GFP-tagged centromere in wild-type and *mps1-R170S* strains. Further, the velocities exhibited by the GFP-tagged centromere as it made poleward migrations were indistinguishable (Fig. 5 D; average forward velocity, *WT* 66.0 nm/sec, n=171, *mps1-R170S* 57.6 nm/sec, n=115; p=0.095; average reverse velocity, *WT* 38.9 nm/sec, n=80, *mps1-R170S* 41.8 nm/sec, n=84; p=0.47, unpaired t tests). If the centromere movements during poleward migration are driven mainly by microtubule depolymerization, then kinetochore microtubule depolymerization occurs at indistinguishable rates in wild-type cells and *mps1-R170S* mutants.

**Figure 5.**
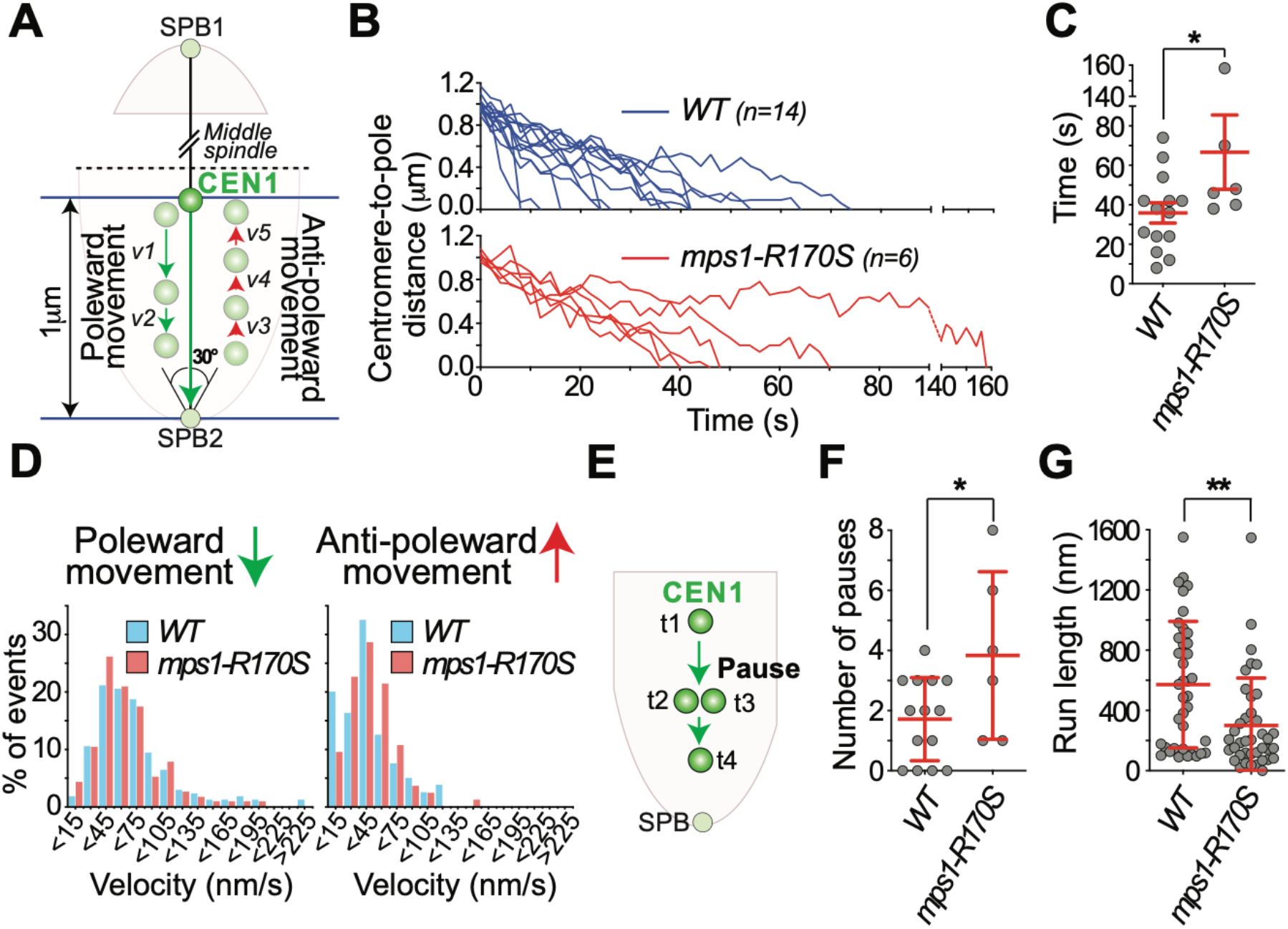
Mps1 is required for processive poleward migration in pro-metaphase. **A.** We identified cells in which the GFP-tagged *CEN1* migrated across the middle of the spindle and proceeded to the opposite pole moving along the central axis of the spindle (within 15° of the axis from the destination SPB). Frame-to-frame movements (both poleward and anti-poleward) for the final one micron of the migration were quantified. **B.** Charts of the poleward movement for wild-type (blue) and *mps1-R170S* mutant (red) cells. T=0 represents the time *CEN1* is one micron from the SPB it is moving toward. **C.** Graph of the cells in (**B**) showing the time spent for each *CEN1* migrating to the SPB from one-micron away. **D.** Distribution of the velocities of the incremental poleward (left) or anti-poleward (right) *CEN1* movements measured during the one-micron poleward migration. **E.** Cartoon illustrating the pauses or reversals of direction of *CEN1* movement observed during the one-micron poleward migrations. **F.** Graph showing the number of pauses or changes of direction for each individual one-micron poleward migration (*WT* 1.7 +/− SD 1.38, n=14; *mps1-R170S*: 3.83+/− SD 2.78, n=6, *p<0.05). **G**. Graphing showing the distance travelled by the GFP-tagged centromere between pauses/reversals (*WT* 571.4 nm +/− SD 420, n=38; *mps1-R170S*: 300.3 nm +/− 315, n=39, **p<0.01). For all graphs, error bars show average and standard deviation, student’s t test was used for statistical comparisons, *p<0.05, **p<0.01.

Since migration to the pole takes much longer in *mps1-R170S* mutants than in wild-type cells but the velocities of poleward movements are indistinguishable, this argues the *mps1-R170S* mutants must pause or reverse more often. To test this, we measured the frequency with which the GFP-tagged centromere paused or reversed direction in its poleward migration (Fig. 5 E). The *MPS1* mutants exhibited significantly more pauses, or reversals of direction, in their journeys to the pole (Fig. 5 F) and the distance travelled between pauses or reversals was significantly shorter (Fig. 5 G). Because the velocities of movement in wild-type cells and *mps1-R170S* mutants are indistinguishable, this results in longer processive movements of the centromere towards the pole in wild-type cells.

If Mps1 acts during prometaphase to promote depolymerization of kinetochore-microtubules, and kinetochore microtubules are stabilized in *MPS1* mutants, then microtubule turnover should be reduced in prometaphase in *MPS1* mutants (Fig. 6 A). To test this, we measured microtubule turnover in cells expressing a photo-convertible mEos2-tagged alpha-tubulin subunit (Markus et al., 2015). mEos2-Tub1 has properties of a green fluorescent protein until it is pulsed with 405 nanometer light, at which point it switches to a red fluorescent protein (Fig. 6 B). To measure turnover of kinetochore microtubules, we pulsed one half of the spindle of cells expressing mEos2-Tub1 with 405 nanometer light, then measured turnover of the red-fluorescent signal (Table 1). Previous measurements of microtubule turnover in budding yeast have been in mitotic cells but the majority of defects we have examined with *MPS1* mutants have been in meiotic cells. Therefore, we first compared microtubule turnover in metaphase spindles of yeast meiotic and mitotic cells and found them to be indistinguishable (Fig. 6 C). To confirm that our methods could detect variations in microtubule turnover rates in meiosis, we measured turnover in cells expressing an auxin-degradable version of the microtubule plus-end protein Stu2 (Stu2-AID*), which helps to regulate microtubule dynamics in mitotic metaphase (Humphrey et al., 2018; Podolski et al., 2014; Miller et al., 2019; 2016; Wolyniak et al., 2006). Cells were induced to enter meiosis and microtubule turnover was measured in the presence or absence of auxin. As observed previously in mitotic cells, (Kosco et al., 2001; Pearson et al., 2003), inactivating Stu2 in meiotic cells reduced microtubule turnover (Fig.6 D). If Mps1 is, like Stu2, promoting microtubule turnover in metaphase cells then inactivating Mps1 should give a similar outcome. To test this, we compared microtubule turnover in metaphase meiotic wild-type cells and *mps1-as1* cells (both in the presence of the Mps1-as1 inhibitor 1-NMPP1). Microtubule turnover rates in metaphase, with or without Mps1 activity, were indistinguishable. This finding is consistent with the reduction in Mps1 levels at kinetochores as they become bi-oriented and the spindle checkpoint is satisfied (Howell et al., 2004; Aravamudhan et al., 2015; Dou et al., 2003; Koch et al., 2019). Our failure to detect a role for Mps1 in metaphase microtubule dynamics could suggest it is simply not involved in that function. The meiotic defects we have observed in *MPS1* mutants were in prometaphase, before chromosomes are bi-oriented, raising the question of whether microtubule dynamics are discernably different in pro-metaphase and metaphase cells using our microtubule turnover assay. In wild-type yeast meiosis, most of the chromosomes are bi-oriented within a few minutes after spindle formation (Meyer et al., 2013). Therefore, we used the *spo11* mutation to obtain a population of cells in which none of the chromosomes are bi-oriented. Consistent with the higher rates of turnover for unattached versus stably attached kinetochore microtubules (Zhai et al., 1995; Gorbsky and Borisy, 1989), the spindles in the *spo11* cells exhibited higher rates of microtubule turnover than were seen in metaphase cells (Fig. 6 B and F). If Mps1, promotes depolymerization of the kinetochore microtubules of non-bi-oriented chromosomes in prometaphase, then this higher rate of turnover seen in prometaphase should be reduced in *MPS1* mutants. For both *mps1-as1* and *mps1-R170S* this proved to be the case (Fig. 6 G and H). Both mutations reduce the rate of turnover to levels like those seen in metaphase cells, where inactivating Mps1 has no discernable effect on microtubule turnover.

**Figure 6.**
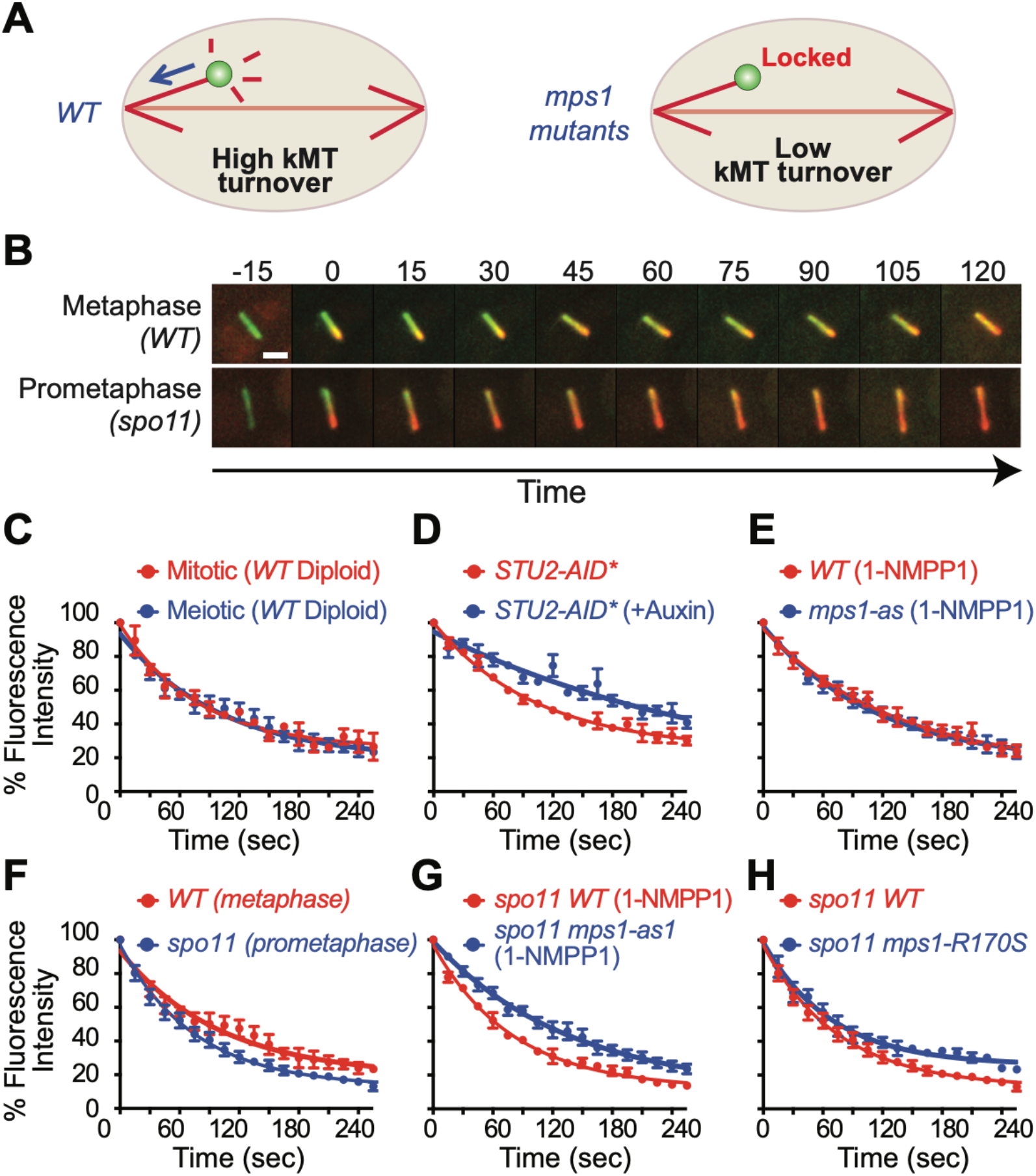
Mps1 promotes microtubule turnover in meiotic prometaphase. **A**. In wild-type cells the shortening kinetochores of actively bi-orienting chromosomes are predicted to cause a high microtubule turnover. *MPS1* mutants exhibit a locked-in-place phenotype that might represent a defect in the depolymerization of kinetochore microtubules. **B.** Cells that were in metaphase or a perpetual prometaphase (*spo11*) were used to measure microtubule turnover. Half-spindles of meiotic cells were pulsed with 405 nm light to photoconvert mEos2-Tub1 (from green to red). Images were acquired every fifteen seconds and the intensity of the red signal was measured (see Methods). Scale bar: 2 μm. **C.** Microtubule turnover on metaphase spindles was measured in a diploid strain undergoing either meiosis or mitosis. **D.** Microtubule turnover was measured in cells expressing STU2-AID* in the presence or absence of auxin and CuSO4 (copper was used to induce expression of the *P_CUP1_-AFB2* F-box protein construct). **E.** Microtubule turnover was measured on meiotic metaphase spindles of wild-type or *mps1-as1* cells in the presence of the Mps1-as1 inhibitor 1-NMPP1. **F.** Microtubule turnover was measured on meiotic metaphase and prometaphase spindles of wild-type cells. **G.** Microtubule turnover was measured on prometaphase spindles (*spo11*) in cells with or without the inactivation of Mps1 by 1-NMPP1. **H.** Microtubule turnover was measured on prometaphase spindles (*spo11*) in wild-type or *mps1-R170S* cells. All experiments show the averages and standard error of the mean of three or more biological replicates with three or more cells per replicate.

## DISCUSSION

Previous work has shown that Mps1 is essential for proper chromosome segregation in meiosis in a variety of organisms (Gilliland et al., 2005; Poss et al., 2004; Straight et al., 2000). We have found that, in budding yeast meiosis, Mps1 is necessary for at least three steps in the bi-orientation process (Meyer et al., 2013; 2018). First, Mps1 is necessary to allow bivalents to migrate to the side-by-side SPBs at the base of a monopolar microtubule array following the exit from meiotic prophase (clustering). Second, Mps1 is necessary for the processive poleward movements on the prometaphase meiosis I spindle that occur before bivalents become bi-oriented. Third, through phosphorylation of Dam1, and possibly other targets, Mps1 stabilizes end-on attachments of the prometaphase kinetochores to microtubules.

The failure of *MPS1* mutants to phosphorylate Dam1 does not explain the massive defects in meiotic chromosome segregation exhibited by *MPS1* mutants. Despite their defects in kMT interactions, *dam1-2A* mutants that cannot be phosphorylated by Mps1 exhibit rather mild meiotic chromosome segregation defects (Shimogawa et al., 2006; Meyer et al., 2018). Thus, there must be another role (or roles) of Mps1 that explains its essentiality for meiotic chromosome segregation. Our experiments have not revealed a critical meiotic substrate but have refined our understanding of the ways in which Mps1 affects chromosome dynamics in meiosis I.

Our results suggest that the major defect in *MPS1* mutants is in regulating microtubule dynamics at the kinetochore interface. A number of observations point to this conclusion. First, the maximum velocity of poleward kinetochore movements in *MPS1* mutants is indistinguishable from that of wild-type cells. This suggests that Mps1 is not required for kinetochores to track de-polymerizing microtubules and is not required for efficient microtubule depolymerization. In *MPS1* mutants, it is rare for kinetochores to traverse the length of the spindle. Instead, kinetochores pause more often than in wild-type cells (Fig. 5 F). These pauses could represent losses of kinetochore microtubule plus-end attachment or pauses in microtubule depolymerization, or both. Given that phosphorylation of Dam1 by Mps1 strengthens kinetochore attachments to plus ends (Shimogawa et al., 2006; Meyer et al., 2018) some of the pauses in *MPS1* mutants are probably due to failures in maintaining the kinetochore-plus-end connection. However, other results suggest that this is not the major defect. First, *MPS1* mutants exhibit low levels of the lagging chromosomes that are an indicator of a defect in attaching kinetochores to microtubules (Meyer et al., 2018; 2013). Second, *MPS1* mutants exhibit a stuck-in-the-middle phenotype in which kinetochores maintain a very stable position in mid-spindle. This is unlike *DAM1* mutants, in which kinetochores and plus-ends become uncoupled, or *NDC80* mutants, in which kinetochores do not attach to microtubules (Meyer et al., 2018). One explanation for the stuck-in-the-middle phenotype is that *MPS1* mutants may be defective in promoting depolymerization of kinetochore-coupled MT plus ends. We propose that when a microtubule plus-end attaches to a kinetochore, the proximity of the microtubule plus-end associated proteins to Mps1 allows Mps1 to phosphorylate key substrates associated with the plus-end, changing their activity or localization in a way that favors microtubule catastrophe over rescue (Fig. 6 A). The identity of these Mps1 substrates and how their phosphorylation biases microtubule dynamics remains an important unanswered question.

The above model does not solve another unknown. Why is it that meiotic chromosome segregation is more vulnerable to defects in Mps1 activity than is mitosis? We offer three possible explanations. First, when mitosis begins, kinetochores are already attached to microtubules. In contrast, the chromosome paring process of meiotic prophase demands that kinetochores be released from microtubules for an extended time period. When meiotic prometaphase begins the kinetochores are dispersed across the nucleus and are then gathered into the microtubule-dense region around the SPBs (clustering) just before the SPBs separate to form a spindle. Mps1 is required for this clustering (Meyer et al., 2013). It may be that in the absence of clustering the formation of initial kinetochore-microtubule attachments on the nascent bipolar spindle is highly inefficient leading to bi-orientation defects. A phenomenon similar to clustering, referred to as kinetochore retrieval, has been reported in *S. pombe* meiosis (Cojoc et al., 2016; Kakui et al., 2013). Here, mutations that lead to defects in meiotic kinetochore retrieval also result in subsequent bi-orientation defects, but it is difficult to know whether the segregation defects are purely due to the failure to cluster the dispersed meiotic kinetochores prior to spindle formation, or to other effects of the mutations.

Second, the vulnerability of meiotic cells to *MPS1* defects might lie in differences between meiotic and mitotic spindles. When yeast meiotic spindles form, most chromosomes are mono-oriented, with most chromosomes clustered near the older SPB (Meyer et al., 2013). Mitosis starts in a similar way (Marco et al., 2013). Thus, in both meiosis and mitosis chromosomes that become bi-oriented have made their way to the spindle mid-zone from the pole. But yeast meiotic spindles are longer, possibly making them more dependent on processes that get them from the poles to the mid-zone (Meyer et al., 2013). Movement from the pole to the mid-zone could be accomplished by pulling of the kinetochore by a long microtubule extending across the spindle from the opposite pole – a process that our results show is defective in *MPS1* mutants (Meyer et al., 2013) both because failure to phosphorylate Dam1 results in defective end-on attachments and because processive poleward movements are defective in *MPS1* mutants. An alternate means to get to the mid-zone from the pole is by movement of chromosomes along microtubules from that pole towards their plus ends. This chromosome gliding mechanism has been reported in in *S. pombe* and animal cells but not budding yeast (Windecker et al., 2009; Akera et al., 2015; Kapoor et al., 2006). In *S. pombe* the process involves proteins (Bub1, Bub3, Mad1, kinesin-5) whose kinetochore localization depend upon Mps1 (Windecker et al., 2009; Akera et al., 2015) and is especially critical for chromosome bi-orientation in cells with long spindles. There is yet no evidence this mechanism is important in budding yeast. However, consistent with this model is the recent demonstration that *BUB1* and *BUB3* mutants, like *MPS1* mutants, both exhibit much higher levels of meiotic than mitotic segregation defects and mis-segregate homologous chromosomes to the older SPB in meiosis I, though not at the high levels seen in *MPS1* mutants (Cairo et al., 2020).

Finally, the flexibility of the connections between homologous meiotic centromeres could make them vulnerable to deficiencies in Mps1. This is true of meiotic chromosomes across species and may explain the shared dependence upon Mps1 in yeast, *Drosophila* and zebrafish meioses. Mitotic sister kinetochores are arranged back-to-back, and tightly cohered. Bi-oriented attachments of sister chromatids are thus probably very quickly under tension and stabilized. In contrast, homologous meiotic kinetochores are connected by chiasmata and therefore a longer tether. This predicts that greater microtubule depolymerization is required in meiosis to separate the homologous kinetochores sufficiently that they are under tension. It may be that in the time interval between the formation of an initial bi-polar attachment, and the generation of stabilizing tension, that one or both of the kinetochore-microtubule connections is lost, and the process must re-start. This more challenging meiotic attachment process may render the cell vulnerable to any defects that diminish the efficiency of establishing kinetochore-microtubule attachments. The observation that in budding yeast, meiotic cells are much more sensitive to defects in the spindle checkpoint than are mitotic cells reinforces the idea that bi-orientation in meiosis faces greater hurdles than in mitosis (Shonn, 2000; Cheslock et al., 2005). But work remains to reveal the greatest vulnerabilities of the meiotic bi-orientation process and how the cell deals with them.

## Abbreviations

SPB: spindle pole body

## ACKNOWLEDGEMENTS

We thank Michael Dresser for providing OMRFQuant Imaging Software and Emma Lee for guidance in using Thru-focus Imaging methods. This project was supported by NIH grant R01GM110271 and OCAST grant HR17-115-3 awarded to DSD and NIH grant R35GM126980 awarded to GJG.

## AUTHOR CONTRIBUTIONS

Conceptualization, REM and DSD; Methodology, REM, ART and DSD; Investigation, REM, ART and DSD; Writing, REM and DSD; Writing – Review and Editing, all authors; Supervision, GJG and DSD; Funding acquisition, GJG and DSD.

## References

Abrieu, A., L. Magnaghi-Jaulin, J.A. Kahana, M. Peter, A. Castro, S. Vigneron, T. Lorca, D.W. Cleveland, and J.C. Labbé. 2001. Mps1 is a kinetochore-associated kinase essential for the vertebrate mitotic checkpoint. Cell. 106:83–93. doi:10.1016/s0092-8674(01)00410-x.

Adams, I.R., and J.V. Kilmartin. 1999. Localization of core spindle pole body (SPB) components during SPB duplication in Saccharomyces cerevisiae. J Cell Biol. 145:809–823.

Akera, T., Y. Goto, M. Sato, M. Yamamoto, and Y. Watanabe. 2015. Mad1 promotes chromosome congression by anchoring a kinesin motor to the kinetochore. Nat Cell Biol. 17:1124–1133. doi:10.1038/ncb3219.

Amberg, D.C., D. Burke, and J.N. Strathern. 2005. Methods in Yeast Genetics. CSHL Press. 1 pp.

Aravamudhan, P., A.A. Goldfarb, and A.P. Joglekar. 2015. The kinetochore encodes a mechanical switch to disrupt spindle assembly checkpoint signalling. Nat. Cell Biol. 17:868–879. doi:10.1038/ncb3179.

Asbury, C.L., D.R. Gestaut, A.F. Powers, A.D. Franck, and T.N. Davis. 2006. The Dam1 kinetochore complex harnesses microtubule dynamics to produce force and movement. Proc Natl Acad Sci USA. 103:9873–9878. doi:10.1073/pnas.0602249103.

Biggins, S., F.F. Severin, N. Bhalla, I. Sassoon, A.A. Hyman, and A.W. Murray. 1999. The conserved protein kinase Ipl1 regulates microtubule binding to kinetochores in budding yeast. Genes Dev. 13:532–544.

Cairo, G., A.M. MacKenzie, and S. Lacefield. 2020. Differential requirement for Bub1 and Bub3 in regulation of meiotic versus mitotic chromosome segregation. J. Cell Biol. 219:1524. doi:10.1083/jcb.201909136.

Cheeseman, I.M., S. Anderson, M. Jwa, E.M. Green, J.S. Kang, J.R. Yates, C.S.M. Chan, D.G. Drubin, and G. Barnes. 2002. Phospho-regulation of kinetochore-microtubule attachments by the Aurora kinase Ipl1p. Cell. 111:163–172.

Chen, J., A. Liao, E.N. Powers, H. Liao, L.A. Kohlstaedt, R. Evans, R.M. Holly, J.K. Kim, M. Jovanovic, and E. Unal. 2020. Aurora B-dependent Ndc80 degradation regulates kinetochore composition in meiosis. Genes Dev. 34:209–225. doi:10.1101/gad.333997.119.

Cheslock, P.S., B.J. Kemp, R.M. Boumil, and D.S. Dawson. 2005. The roles of MAD1, MAD2 and MAD3 in meiotic progression and the segregation of nonexchange chromosomes. Nat Genet. 37:756–760. doi:10.1038/ng1588.

Chmátal, L., K. Yang, R.M. Schultz, and M.A. Lampson. 2015. Spatial Regulation of Kinetochore Microtubule Attachments by Destabilization at Spindle Poles in Meiosis I. Curr. Biol. 25:1835–1841. doi:10.1016/j.cub.2015.05.013.

Cojoc, G., A.-M. Florescu, A. Krull, A.H. Klemm, N. Pavin, F. Jülicher, and I.M. Tolić. 2016. Paired arrangement of kinetochores together with microtubule pivoting and dynamics drive kinetochore capture in meiosis I. Sci Rep. 6:25736–12. doi:10.1038/srep25736.

Conrad, M.N., C.-Y. Lee, G. Chao, M. Shinohara, H. Kosaka, A. Shinohara, J.-A. Conchello, and M.E. Dresser. 2008. Rapid Telomere Movement in Meiotic Prophase Is Promoted By NDJ1, MPS3, and CSM4 and Is Modulated by Recombination. Cell. 133:1175–1187. doi:10.1016/j.cell.2008.04.047.

Daum, J.R., J.D. Wren, J.J. Daniel, S. Sivakumar, J.N. McAvoy, T.A. Potapova, and G.J. Gorbsky. 2009. Ska3 is required for spindle checkpoint silencing and the maintenance of chromosome cohesion in mitosis. Curr. Biol. 19:1467–1472. doi:10.1016/j.cub.2009.07.017.

Dou, Z., A. Sawagechi, J. Zhang, H. Luo, L. Brako, and X.B. Yao. 2003. Dynamic distribution of TTK in HeLa cells: insights from an ultrastructural study. Cell Res. 13:443–449. doi:10.1038/sj.cr.7290186.

Dresser, M.E., D.J. Ewing, S.N. Harwell, D. Coody, and M.N. Conrad. 1994. Nonhomologous synapsis and reduced crossing over in a heterozygous paracentric inversion in Saccharomyces cerevisiae. Genetics. 138:633–647.

Franco, A., J.C. Meadows, and J.B.A. Millar. 2007. The Dam1/DASH complex is required for the retrieval of unclustered kinetochores in fission yeast. J Cell Sci. 120:3345–3351. doi:10.1242/jcs.013698.

Gachet, Y., C. Reyes, T. Courthéoux, S. Goldstone, G. Gay, C. Serrurier, and S. Tournier. 2008. Sister kinetochore recapture in fission yeast occurs by two distinct mechanisms, both requiring Dam1 and Klp2. Mol. Biol. Cell. 19:1646–1662. doi:10.1091/mbc.E07-09-0910.

Gaitanos, T.N., A. Santamaria, A.A. Jeyaprakash, Bin Wang, E. Conti, and E.A. Nigg. 2009. Stable kinetochore-microtubule interactions depend on the Ska complex and its new component Ska3/C13Orf3. The EMBO Journal. 28:1442–1452. doi:10.1038/emboj.2009.96.

Gilliland, W.D., S.M. Wayson, and R.S. Hawley. 2005. The meiotic defects of mutants in the Drosophila mps1 gene reveal a critical role of Mps1 in the segregation of achiasmate homologs. Curr Biol. 15:672–677. doi:10.1016/j.cub.2005.02.062.

Godek, K.M., L. Kabeche, and D.A. Compton. 2015. Regulation of kinetochore-microtubule attachments through homeostatic control during mitosis. Nat Rev Mol Cell Biol. 16:57–64. doi:10.1038/nrm3916.

Gorbsky, G.J., and G.G. Borisy. 1989. Microtubules of the kinetochore fiber turn over in metaphase but not in anaphase. J Cell Biol. 109:653–662. doi:10.1083/jcb.109.2.653.

Grishchuk, E.L., A.K. Efremov, V.A. Volkov, I.S. Spiridonov, N. Gudimchuk, S. Westermann, D. Drubin, G. Barnes, J.R. McIntosh, and F.I. Ataullakhanov. 2008. The Dam1 ring binds microtubules strongly enough to be a processive as well as energy-efficient coupler for chromosome motion. Proceedings of the National Academy of Sciences. 105:15423–15428. doi:10.1073/pnas.0807859105.

Hardwick, K.G., E. Weiss, F.C. Luca, M. Winey, and A.W. Murray. 1996. Activation of the budding yeast spindle assembly checkpoint without mitotic spindle disruption. Science. 273:953–956.

Hayden, J.H., S.S. Bowser, and C.L. Rieder. 1990. Kinetochores capture astral microtubules during chromosome attachment to the mitotic spindle: direct visualization in live newt lung cells. J Cell Biol. 111:1039–1045. doi:10.1083/jcb.111.3.1039.

Howell, B.J., B. Moree, E.M. Farrar, S. Stewart, G. Fang, and E.D. Salmon. 2004. Spindle checkpoint protein dynamics at kinetochores in living cells. Curr Biol. 14:953–964. doi:10.1016/j.cub.2004.05.053.

Humphrey, L., I. Felzer-Kim, and A.P. Joglekar. 2018. Stu2 acts as a microtubule destabilizer in metaphase budding yeast spindles. Mol. Biol. Cell. 29:247–255. doi:10.1091/mbc.E17-08-0494.

Janke, C., M.M. Magiera, N. Rathfelder, C. Taxis, S. Reber, H. Maekawa, A. Moreno-Borchart, G. Doenges, E. Schwob, E. Schiebel, and M. Knop. 2004. A versatile toolbox for PCR-based tagging of yeast genes: new fluorescent proteins, more markers and promoter substitution cassettes. Yeast. 21:947–962. doi:10.1002/yea.1142.

Jones, M.H., B.J. Huneycutt, C.G. Pearson, C. Zhang, G. Morgan, K. Shokat, K. Bloom, and M. Winey. 2005. Chemical genetics reveals a role for Mps1 kinase in kinetochore attachment during mitosis. Curr Biol. 15:160–165. doi:10.1016/j.cub.2005.01.010.

Kakui, Y., M. Sato, N. Okada, T. Toda, and M. Yamamoto. 2013. Microtubules and Alp7-Alp14 (TACC-TOG) reposition chromosomes before meiotic segregation. Nat Cell Biol. 15:786–796. doi:10.1038/ncb2782.

Kapoor, T.M., M.A. Lampson, P. Hergert, L. Cameron, D. Cimini, E.D. Salmon, B.F. McEwen, and A. Khodjakov. 2006. Chromosomes can congress to the metaphase plate before biorientation. Science. 311:388–391. doi:10.1126/science.1122142.

Kitamura, E., K. Tanaka, Y. Kitamura, and T.U. Tanaka. 2007. Kinetochore microtubule interaction during S phase in Saccharomyces cerevisiae. Genes Dev. 21:3319–3330. doi:10.1101/gad.449407.

Klapholz, S., C.S. Waddell, and R.E. Esposito. 1985. The role of the SPO11 gene in meiotic recombination in yeast. Genetics. 110:187–216.

Koch, L.B., K.N. Opoku, Y. Deng, A. Barber, A.J. Littleton, N. London, S. Biggins, and C.L. Asbury. 2019. Autophosphorylation is sufficient to release Mps1 kinase from native kinetochores. Proceedings of the National Academy of Sciences. 116:17355–17360. doi:10.1073/pnas.1901653116.

Kosco, K.A., C.G. Pearson, P.S. Maddox, P.J. Wang, I.R. Adams, E.D. Salmon, K. Bloom, and T.C. Huffaker. 2001. Control of microtubule dynamics by Stu2p is essential for spindle orientation and metaphase chromosome alignment in yeast. Mol Biol Cell. 12:2870–2880. doi:10.1091/mbc.12.9.2870.

Koszul, R., K.P. Kim, M. Prentiss, N. Kleckner, and S. Kameoka. 2008. Meiotic chromosomes move by linkage to dynamic actin cables with transduction of force through the nuclear envelope. Cell. 133:1188–1201. doi:10.1016/j.cell.2008.04.050.

Lampert, F., P. Hornung, and S. Westermann. 2010. The Dam1 complex confers microtubule plus end-tracking activity to the Ndc80 kinetochore complex. J. Cell Biol. 189:641–649. doi:10.1083/jcb.200912021.

Lampson, M.A., and E.L. Grishchuk. 2017. Mechanisms to Avoid and Correct Erroneous Kinetochore-Microtubule Attachments. Biology 2017, Vol. 6, Page 12. 6:1. doi:10.3390/biology6010001.

Loidl, J., F. Klein, and H. Scherthan. 1994. Homologous pairing is reduced but not abolished in asynaptic mutants of yeast. J Cell Biol. 125:1191–1200.

Longtine, M.S., A. McKenzie, D.J. Demarini, N.G. Shah, A. Wach, A. Brachat, P. Philippsen, and J.R. Pringle. 1998. Additional modules for versatile and economical PCR-based gene deletion and modification in Saccharomyces cerevisiae. Yeast. 14:953–961. doi:10.1002/(SICI)1097-0061(199807)14:10<953::AID-YEA293>3.0.CO;2-U.

Magidson, V., C.B. O’Connell, J. Lončarek, R. Paul, A. Mogilner, and A. Khodjakov. 2011. The spatial arrangement of chromosomes during prometaphase facilitates spindle assembly. Cell. 146:555–567. doi:10.1016/j.cell.2011.07.012.

Marco, E., J.F. Dorn, P.-H. Hsu, K. Jaqaman, P.K. Sorger, and G. Danuser. 2013. S. cerevisiae chromosomes biorient via gradual resolution of syntely between S phase and anaphase. Cell. 154:1127–1139. doi:10.1016/j.cell.2013.08.008.

Markus, S.M., S. Omer, K. Baranowski, and W.-L. Lee. 2015. Improved Plasmids for Fluorescent Protein Tagging of Microtubules in Saccharomyces cerevisiae. Traffic. 16:773–786. doi:10.1111/tra.12276.

Merdes, A., and J. De Mey. 1990. The mechanism of kinetochore-spindle attachment and polewards movement analyzed in PtK2 cells at the prophase-prometaphase transition. Eur. J. Cell Biol. 53:313–325.

Meyer, R.E., H.H. Chuong, M. Hild, C.L. Hansen, M. Kinter, and D.S. Dawson. 2015. Ipl1/Aurora-B is necessary for kinetochore restructuring in meiosis I in Saccharomyces cerevisiae. Mol. Biol. Cell. 26:2986–3000. doi:10.1091/mbc.E15-01-0032.

Meyer, R.E., J. Brown, L. Beck, and D.S. Dawson. 2018. Mps1 promotes chromosome meiotic chromosome biorientation through Dam1. Mol. Biol. Cell. 29:479–489. doi:10.1091/mbc.E17-08-0503.

Meyer, R.E., S. Kim, D. Obeso, P.D. Straight, M. Winey, and D.S. Dawson. 2013. Mps1 and Ipl1/Aurora B act sequentially to correctly orient chromosomes on the meiotic spindle of budding yeast. Science. 339:1071–1074. doi:10.1126/science.1232518.

Miller, M.P., C.L. Asbury, and S. Biggins. 2016. A TOG Protein Confers Tension Sensitivity to Kinetochore-Microtubule Attachments. Cell. 165:1428–1439. doi:10.1016/j.cell.2016.04.030.

Miller, M.P., E. Unal, G.A. Brar, and A. Amon. 2012. Meiosis I chromosome segregation is established through regulation of microtubule-kinetochore interactions. eLife. 1:e00117–e00117. doi:10.7554/eLife.00117.057.

Miller, M.P., R.K. Evans, A. Zelter, E.A. Geyer, M.J. MacCoss, L.M. Rice, T.N. Davis, C.L. Asbury, and S. Biggins. 2019. Kinetochore-associated Stu2 promotes chromosome biorientation in vivo. PLoS Genet. 15:e1008423. doi:10.1371/journal.pgen.1008423.

Monje-Casas, F., V.R. Prabhu, B.H. Lee, M. Boselli, and A. Amon. 2007. Kinetochore orientation during meiosis is controlled by Aurora B and the monopolin complex. Cell. 128:477–490. doi:10.1016/j.cell.2006.12.040.

Nicklas, R.B. 1997. How cells get the right chromosomes. Science. 275:632–637.

Pearson, C.G., P.S. Maddox, T.R. Zarzar, E.D. Salmon, and K. Bloom. 2003. Yeast kinetochores do not stabilize Stu2p-dependent spindle microtubule dynamics. Mol Biol Cell. 14:4181–4195. doi:10.1091/mbc.E03-03-0180.

Podolski, M., M. Mahamdeh, and J. Howard. 2014. Stu2, the budding yeast XMAP215/Dis1 homolog, promotes assembly of yeast microtubules by increasing growth rate and decreasing catastrophe frequency. Journal of Biological Chemistry. 289:28087–28093. doi:10.1074/jbc.M114.584300.

Poss, K.D., A. Nechiporuk, K.F. Stringer, C. Lee, and M.T. Keating. 2004. Germ cell aneuploidy in zebrafish with mutations in the mitotic checkpoint gene mps1. Genes Dev. 18:1527–1532. doi:10.1101/gad.1182604.

Powers, A.F., A.D. Franck, D.R. Gestaut, J. Cooper, B. Gracyzk, R.R. Wei, L. Wordeman, T.N. Davis, and C.L. Asbury. 2009. The Ndc80 kinetochore complex forms load-bearing attachments to dynamic microtubule tips via biased diffusion. Cell. 136:865–875. doi:10.1016/j.cell.2008.12.045.

Rieder, C.L., and S.P. Alexander. 1990. Kinetochores are transported poleward along a single astral microtubule during chromosome attachment to the spindle in newt lung cells. J Cell Biol. 110:81–95. doi:10.1083/jcb.110.1.81.

Sarangapani, K.K., E. Duro, Y. Deng, F. de L. Alves, Q. Ye, K.N. Opoku, S. Ceto, J. Rappsilber, K.D. Corbett, S. Biggins, A.L. Marston, and C.L. Asbury. 2014. Sister kinetochores are mechanically fused during meiosis I in yeast. Science. 346:248–251. doi:10.1126/science.1256729.

Schmidt, J.C., H. Arthanari, A. Boeszoermenyi, N.M. Dashkevich, E.M. Wilson-Kubalek, N. Monnier, M. Markus, M. Oberer, R.A. Milligan, M. Bathe, G. Wagner, E.L. Grishchuk, and I.M. Cheeseman. 2012. The kinetochore-bound Ska1 complex tracks depolymerizing microtubules and binds to curved protofilaments. Dev Cell. 23:968–980. doi:10.1016/j.devcel.2012.09.012.

Shimogawa, M.M., B. Graczyk, M.K. Gardner, S.E. Francis, E.A. White, M. Ess, J.N. Molk, C. Ruse, S. Niessen, J.R. Yates, E.G.D. Muller, K. Bloom, D.J. Odde, and T.N. Davis. 2006. Mps1 phosphorylation of Dam1 couples kinetochores to microtubule plus ends at metaphase. Curr Biol. 16:1489–1501. doi:10.1016/j.cub.2006.06.063.

Shonn, M.A. 2000. Requirement of the Spindle Checkpoint for Proper Chromosome Segregation in Budding Yeast Meiosis. 289:300–303. doi:10.1126/science.289.5477.300.

Straight, A.F., A.S. Belmont, C.C. Robinett, and A.W. Murray. 1996. GFP tagging of budding yeast chromosomes reveals that protein-protein interactions can mediate sister chromatid cohesion. Curr Biol. 6:1599–1608.

Straight, P.D., T.H. Giddings, and M. Winey. 2000. Mps1p regulates meiotic spindle pole body duplication in addition to having novel roles during sporulation. Mol Biol Cell. 11:3525–3537. doi:10.1091/mbc.11.10.3525.

Tanaka, K., E. Kitamura, Y. Kitamura, and T.U. Tanaka. 2007. Molecular mechanisms of microtubule-dependent kinetochore transport toward spindle poles. J. Cell Biol. 178:269–281. doi:10.1083/jcb.200702141.

Tanaka, K., N. Mukae, H. Dewar, M. van Breugel, E.K. James, A.R. Prescott, C. Antony, and T.U. Tanaka. 2005. Molecular mechanisms of kinetochore capture by spindle microtubules. Nature. 434:987–994. doi:10.1038/nature03483.

Tanaka, T.U. 2010. Kinetochore-microtubule interactions: steps towards bi-orientation. EMBO J. 29:4070–4082. doi:10.1038/emboj.2010.294.

Tanaka, T.U., N. Rachidi, C. Janke, G. Pereira, M. Galova, E. Schiebel, M.J.R. Stark, and K. Nasmyth. 2002. Evidence that the Ipl1-Sli15 (Aurora kinase-INCENP) complex promotes chromosome bi-orientation by altering kinetochore-spindle pole connections. Cell. 108:317–329.

Umbreit, N.T., M.P. Miller, J.F. Tien, J. Cattin Ortolá, L. Gui, K.K. Lee, S. Biggins, C.L. Asbury, and T.N. Davis. 2014. Kinetochores require oligomerization of Dam1 complex to maintain microtubule attachments against tension and promote biorientation. Nat Commun. 5:4951. doi:10.1038/ncomms5951.

Volkov, V.A., A.V. Zaytsev, N. Gudimchuk, P.M. Grissom, A.L. Gintsburg, F.I. Ataullakhanov, J.R. McIntosh, and E.L. Grishchuk. 2013. Long tethers provide high-force coupling of the Dam1 ring to shortening microtubules. Proceedings of the National Academy of Sciences. 110:7708–7713. doi:10.1073/pnas.1305821110.

Weiss, E., and M. Winey. 1996. The Saccharomyces cerevisiae spindle pole body duplication gene MPS1 is part of a mitotic checkpoint. J Cell Biol. 132:111–123. doi:10.1083/jcb.132.1.111.

Welburn, J.P.I., E.L. Grishchuk, C.B. Backer, E.M. Wilson-Kubalek, J.R. Yates, and I.M. Cheeseman. 2009. The human kinetochore Ska1 complex facilitates microtubule depolymerization-coupled motility. Dev Cell. 16:374–385. doi:10.1016/j.devcel.2009.01.011.

Westermann, S., H.-W. Wang, A. Avila-Sakar, D.G. Drubin, E. Nogales, and G. Barnes. 2006. The Dam1 kinetochore ring complex moves processively on depolymerizing microtubule ends. Nature. 440:565–569. doi:10.1038/nature04409.

Windecker, H., M. Langegger, S. Heinrich, and S. Hauf. 2009. Bub1 and Bub3 promote the conversion from monopolar to bipolar chromosome attachment independently of shugoshin. EMBO Rep. 10:1022–1028. doi:10.1038/embor.2009.183.

Winey, M., G.P. Morgan, P.D. Straight, T.H. Giddings, and D.N. Mastronarde. 2005. Three-dimensional ultrastructure of Saccharomyces cerevisiae meiotic spindles. Mol Biol Cell. 16:1178–1188. doi:10.1091/mbc.E04-09-0765.

Wolyniak, M.J., K. Blake-Hodek, K. Kosco, E. Hwang, L. You, and T.C. Huffaker. 2006. The regulation of microtubule dynamics in Saccharomyces cerevisiae by three interacting plus-end tracking proteins. Mol Biol Cell. 17:2789–2798. doi:10.1091/mbc.E05-09-0892.

Zhai, Y., P.J. Kronebusch, and G.G. Borisy. 1995. Kinetochore microtubule dynamics and the metaphase-anaphase transition. J Cell Biol. 131:721–734. doi:10.1083/jcb.131.3.721.

