## Supplemental Figure and Tables for "Mps1 promotes poleward chromosome movements in meiotic pro-metaphase"

**SUPPLEMENTAL MATERIALS (Meyer et al.)**

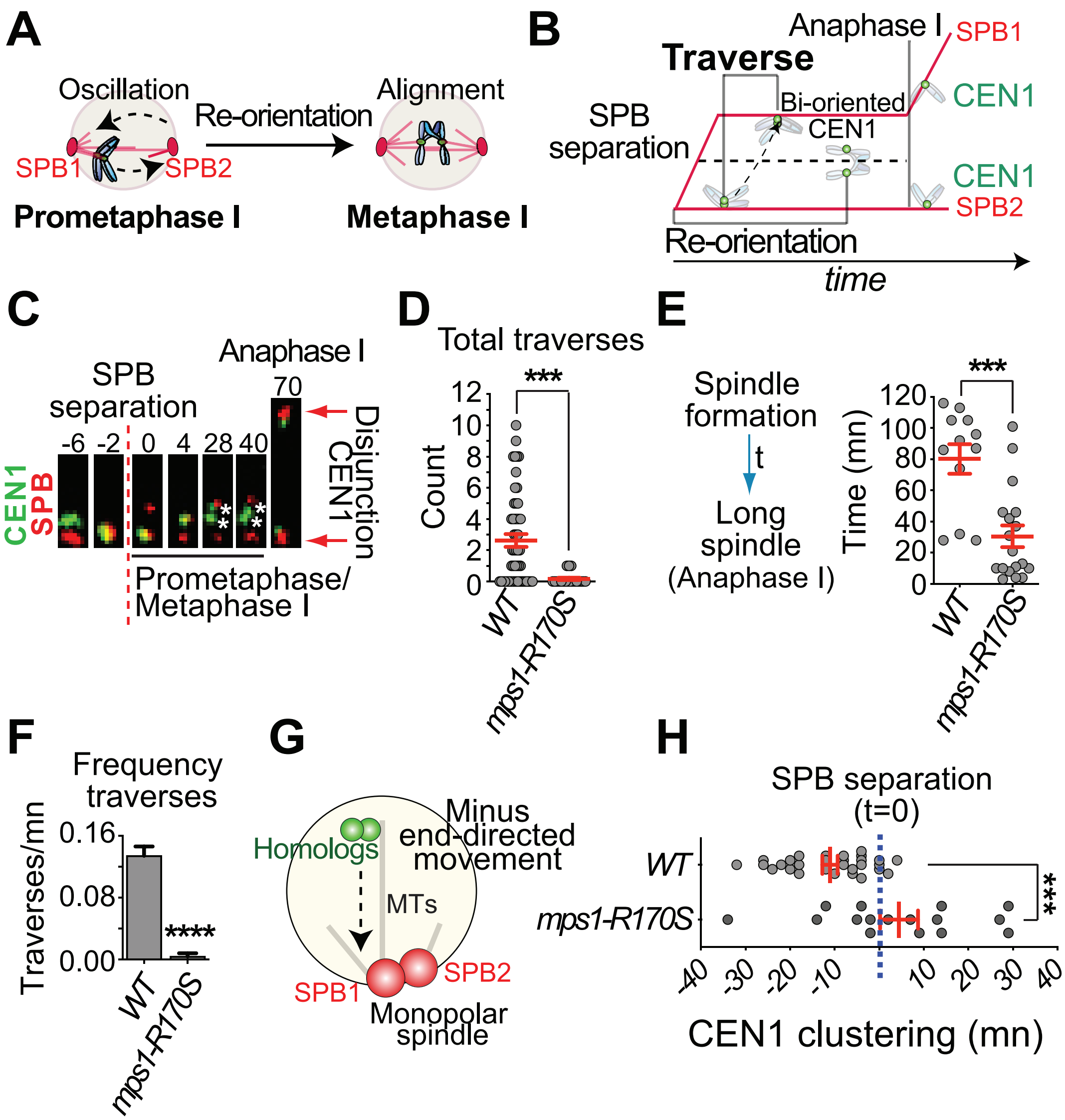


**Figure S1. Mps1 is required for the bi-orientation of bivalents in meiosis. A.** Cartoon illustrating the process of re-orientation and chromosome alignment. The vast majority of chromosome pairs initially attach to one spindle pole in early prometaphase I, thus the re-orientation process is a necessary step to promote the establishment of bi-oriented attachments for each chromosome pair. **B.** The re-orientation process can be quantified by counting the number of traverses (movements across the entire spindle).

For simplicity only *CEN1* is represented. **C-H** Strains evaluated were all diploids carrying GFP-tagged centromeres of chromosome *I* (CEN1-GFP*)* and expressing *SPC42-DsRed* to mark the SPBs. Cells were sporulated and released from a pachytene arrest ($P_{GAL1}$*-NDT80, GAL4-ER*) at 6 hours after meiotic induction by the addition of 5 µM β-estradiol. The diploid cells were observed by time-lapse imaging during meiosis at two-minute intervals for 3-4 hours. **C.** Representative cell from wild-type is shown. The asterisks indicate split homologous centromeres. Diploid cells (wild-type and *mps1-R170S* mutants) that were able to form bivalents (*SPO11*) were analyzed for the following parameters: **D.** The total number of traverses until bi-orientation or until anaphase I (in cases where bi-orientation of chromosome *I* was not achieved) was determined for each cell. **E.** The duration of metaphase I was determined for each individual cell (n≥18) by calculating the time between SPB separation and the formation of a long bipolar spindle (≥3.5 µm). The shortened metaphase in *mps1-R170S* mutants highlights their loss of the spindle assembly checkpoint. **F.** The frequency of *CEN1* traverses per minute during the first twenty minutes of prometaphase (marked by the separation of SPBs) is represented. **G.** Schematic representation of the process of chromosome movement on a monopolar spindle which occurs before bipolar spindle formation. **H.** The timing of the final *CEN1* clustering (when CEN1 reaches the SPB and remains associated with it) was monitored relative to the moment of SPB separation for each individual cell of the indicated genotypes (n≥19). The red dotted line represents the time of SPB separation. ***$p < 0.001$, ****$p < 0.0001$ (student's t test).

### <u>Table S1: Diploid strain list</u>

| Diploid | Parents strain | Figure |
|---|---|---|
| DRM3032 | X1151 * Y1114 | Fig. 2, 3 "*WT*" |
| DRM2417 | X1195 * Y1144 | Fig. 2, 3 "*mps1-R170S*" |
| DRM1658 | X1195 * Y1147 | Fig. 2, 3 "*mps1-as1*" |
| DRM1661 | X1197 * Y1148 | Fig. 2, 3 "*mps1-as1*" |
| DRM3107 | X2297 * Y2145 | Fig. 3 "*ndc80-md*" |
| DRM2549 | X1151 * Y2083 | Fig. 4, 5 "*WT*" |
| DRM2550 | X1152 * Y2083 | Fig. 4, 5 "*WT*" |
| DRM2525 | X1196 * Y2099 | Fig. 4, 5 "*mps1-R170S*" |
| DRM2526 | X1197 * Y2100 | Fig. 4, 5 "*mps1-R170S*" |
| DRM2551 | X1196 * Y2091 | Fig. 4, 5 "*mps1-R170S*" |
| DRM2552 | X1197 * Y2092 | Fig. 4, 5 "*mps1-R170S*" |
| DRM2649 | X2297 * Y2187 | Fig. 4 "*ndc80-md*" |
| DRM2650 | X2298 * Y2188 | Fig. 4 "*ndc80-md*" |
| DRM3558/ DRM3784 | X3217 * Y2771 | Fig. 6 "Mitotic *(WT* Diploid*)"* |
| <u>DRM3451</u> | X3227 * Y2779 | Fig. 6, S2B "Meiotic *(WT* Diploid*)", "WT (1-NMPP1)", "WT (metaphase)"* |
| <u>DRM3452</u> | X3228 * Y2780 | Fig. 6, S2B "Meiotic *(WT* Diploid*)", "WT (1-NMPP1)", "WT (metaphase)"* |
| DRM3455 | X3233 * Y2783 | Fig. 6, S2B "*mps1-as1 (1-NMPP1)"* |
| DRM3456 | X3234 * Y2784 | Fig. 6, S2B "*mps1-as1 (1-NMPP1)"* |
| DRM3457 | X3231 * Y2781 | Fig. 6, S2B "*spo11 (prometaphase)", "spo11 WT (1-NMPP1)", "spo11 WT"* |
| DRM3458 | X3232 * Y2782 | Fig. 6, S2B "*spo11 (prometaphase)", "spo11 WT (1-NMPP1)", "spo11 WT",* |
| DRM3459 | X3229 * Y2787 | Fig. 6, S2B "*spo11 mps1-R170S"* |
| DRM3460 | X3230 * Y2787 | Fig. 6, S2B "*spo11 mps1-R170S"* |
| DRM3479 | X3229 * Y2805 | Fig. 6, S2B "*spo11 mps1-as1 (1-NMPP1)"* |
| DRM3480 | X3230 * Y2805 | Fig. 6, S2B "*spo11 mps1-as1 (1-NMPP1)"* |
| DRM3893 | X3383 * Y2933 | Fig. 6, S2B "*STU2-AID*", "STU2-AID* (+Auxin)"* |
| DRM3894 | X3384 * Y2934 | Fig. 6, S2B "*STU2-AID*", "STU2-AID* (+Auxin)"* |
| DRM2405 | X1843 * Y1693 | Fig. S1 "*WT*" |
| DRM2415 | X1142 * Y1136 | Fig. S1 "*mps1-R170S*" |
| DRM1967 | X1103 * Y1092 | Fig. S2A "*WT*" |
| DRM1969 | X1142 * Y1124 | Fig. S2A "*mps1-as1*" |

### <u>Table S2: Haploid strain list</u>

| | |
|---|---|
| X1843 | *MATa, trp1-Δ63, his3-Δ1, leu2, met13-d, tyr1-1, lys2::pLL1[PCYC1-GFP-lacI LYS2], can1-R, ura3::pKB80 [PGPD1-GAL4(848)-ER-URA3::hphNT1], natNT2-PGAL1-NDT80, CEN1::pJN2[lacO256 LEU2]* |
| Y1693 | *MAT∝, ura3::pAFS152[URA3 PCYC-GFP-lacI], trp1-Δ63, his3-Δ1, leu2, lys2::pMDE798[PDMC1-GFP-lacI], tyr1-2, met13-c, cyh2-1, SPC42-[MDE1145: URA3 SPC42-DSRed], natNT2-PGAL1-NDT80, CEN1::pJN2[lacO256 LEU2]* |
| X1142 | *MATa, ura3::pKB80 [PGPD1-GAL4(848)-ER-URA3::hphNT1], trp1-63, his3-1, leu2, met13-d, tyr1-1, lys2::pLL1[PCYC1-GFP-lacI LYS2], can1 ®, ZIP1-TEV?, KanMX-PGAL1-NDT80, PCLB2-3HA-MPS1 KANMX6, CEN1::pJN2[lacO256 LEU2]* |
| Y1136 | *MAT∝, ura3::pAFS152[URA3 PCYC-GFP-lacI], trp1-Δ63, his3-Δ1, leu2, lys2::pMDE798[PDMC1-GFP-lacI], tyr1-2, met13-c, cyh2-1, SPC42-[MDE1145: URA3 SPC42-DSRed], KanMX-PGAL1-NDT80, mps1-R170S::his5, CEN1::pJN2[lacO256 LEU2]* |
| X1151 | *MAT∝, ura3::pKB80 [PGPD1-GAL4(848)-ER-URA3::hphNT1], trp1-63, his3-1, leu2, met13-d, tyr1-1, lys2::pLL1[PCYC1-GFP-lacI LYS2], can1 ®, ZIP1-TEV?, KanMX-PGAL1-NDT80, CEN1::pJN2[lacO256 LEU2], spo11::KANMX* |
| Y1114 | *MATa, ura3::pAFS152[URA3 PCYC-GFP-lacI], trp1-Δ63, his3-Δ1, leu2, lys2::pMDE798[PDMC1-GFP-lacI], tyr1-2, met13-c, cyh2-1, SPC42-[MDE1145: URA3 SPC42-DSRed], KanMX-PGAL1-NDT80, spo11::KANMX* |
| X1195 | *MATa, ura3::pKB80 [PGPD1-GAL4(848)-ER-URA3::hphNT1], trp1-63, his3-1, leu2, met13-d, tyr1-1, lys2::pLL1[PCYC1-GFP-lacI LYS2], can1 ®, ZIP1-TEV?, KanMX-PGAL1-NDT80, spo11::KANMX, PCLB2-3HA-MPS1 KANMX6, CEN1::pJN2[lacO256 LEU2]* |
| Y1144 | *MAT∝, ura3::pAFS152[URA3 PCYC-GFP-lacI], trp1-Δ63, his3-Δ1, leu2, lys2::pMDE798[PDMC1-GFP-lacI], tyr1-2, met13-c, cyh2-1, SPC42-[MDE1145: URA3 SPC42-DSRed], KanMX-PGAL1-NDT80, spo11::KANMX, mps1-R170S::his5* |
| Y1147 | MAT∝, ura3::pAFS152[URA3 PCYC-GFP-lacI], trp1-Δ63, his3-Δ1, leu2, lys2::pMDE798[PDMC1-GFP-lacI], tyr1-2, met13-c, cyh2-1, SPC42-[MDE1145: URA3 SPC42-DSRed], KanMX-PGAL1-NDT80, spo11::KANMX, mps1Δ::KANMX, TRP1::10Xmyc-mps1-as1 |
| X1197 | *MAT∝, ura3::pKB80 [$P_{GPD1}$-GAL4(848)-ER-URA3::hphNT1], trp1-63, his3-1, leu2, met13-d, tyr1-1, lys2::pLL1[PCYC1-GFP-lacI LYS2], can1 ®, ZIP1-TEV?, KanMX-$P_{GAL1}$-NDT80, spo11::KANMX, PCLB2-3HA-MPS1 KANMX6, CEN1::pJN2[lacO256 LEU2]* |
| Y1148 | *MATa, ura3::pAFS152[URA3 PCYC-GFP-lacI], trp1-Δ63, his3-Δ1, leu2, lys2::pMDE798[PDMC1-GFP-lacI], tyr1-2, met13-c, cyh2-1, SPC42-[MDE1145: URA3 SPC42-DSRed], KanMX-PGAL1-NDT80, spo11::KANMX, mps1Δ::KANMX, TRP1::10Xmyc-mps1-as1* |
| X2297 | *MATa, trp1-Δ63, his3-Δ1, leu2, met13-d, tyr1-1, lys2::pLL1[PCYC1-GFP-lacI LYS2], can1-R, ura3::pKB80 [PGPD1-GAL4(848)-ER-URA3::hphNT1], natNT2-PGAL1-NDT80, CEN1::pJN2[lacO256 LEU2], PCLB2-3HA-NDC80 KanMX6, spo11::HIS3MX6* |
| Y2145 | *MAT∝, ura3::pAFS152[URA3 PCYC-GFP-lacI], trp1-Δ63, his3-Δ1, leu2, lys2::pMDE798[PDMC1-GFP-lacI], tyr1-2, met13-c, cyh2-1, SPC42-[MDE1145: URA3 SPC42-DSRed], natNT2-PGAL1-NDT80, PCLB2-3HA-NDC80 KanMX6, spo11::HIS3MX6* |
| Y2083 | *MATa, trp1-Δ63, his3-Δ1, leu2, lys2::pLL1[PCYC1-GFP-lacI LYS2], tyr1-2, met13-c, cyh2-1, ura3::pAFS152[URA3 PCYC1-GFP-lacI], SPC42-GFP-TRP1 , natNT2-PGAL1-NDT80, spo11::KANMX* |
| X1152 | *MAT∝, ura3::pKB80 [PGPD1-GAL4(848)-ER-URA3::hphNT1], trp1-63, his3-1, leu2, met13-d, tyr1-1, lys2::pLL1[PCYC1-GFP-lacI LYS2], can1 ®, ZIP1-TEV?, KanMX-PGAL1-NDT80, CEN1::pJN2[lacO256 LEU2], spo11::KANMX* |

| | |
|---|---|
| X1196 | MATa, ura3::pKB80 [$P_{GPD1}$-GAL4(848)-ER-URA3::hphNT1], trp1-63, his3-1, leu2, met13-d, tyr1-1, lys2::pLL1[PCYC1-GFP-lacI LYS2], can1 ®, ZIP1-TEV?, KanMX-$P_{GAL1}$-NDT80, spo11::KANMX, PCLB2-3HA-MPS1 KANMX6, CEN1::pJN2[lacO256 LEU2] |
| Y2099 | MAT∝, ura3::pAFS152[URA3 PCYC-GFP-lacI], trp1-Δ63, his3-Δ1, leu2, lys2::pMDE798[PDMC1-GFP-lacI], tyr1-2, met13-c, cyh2-1, SPC42-GFP-TRP1, spo11::HIS3MX6, natNT2-PGAL1-NDT80, mps1-R170S::his5 |
| Y2100 | MATa, ura3::pAFS152[URA3 PCYC-GFP-lacI], trp1-Δ63, his3-Δ1, leu2, lys2::pMDE798[PDMC1-GFP-lacI], tyr1-2, met13-c, cyh2-1, SPC42-GFP-TRP1, spo11::HIS3MX6, natNT2-PGAL1-NDT80, mps1-R170S::his5 |
| Y2091 | MAT∝, trp1-Δ63, his3-Δ1, leu2, lys2::pLL1[PCYC1-GFP-lacI LYS2], tyr1-2, met13-c, cyh2-1, ura3::pAFS152[URA3 PCYC1-GFP-lacI], SPC42-GFP-TRP1 , natNT2-PGAL1-NDT80, spo11::KANMX, mps1-R170S::his5 |
| Y2092 | MATa, trp1-Δ63, his3-Δ1, leu2, lys2::pLL1[PCYC1-GFP-lacI LYS2], tyr1-2, met13-c, cyh2-1, ura3::pAFS152[URA3 PCYC1-GFP-lacI], SPC42-GFP-TRP1 , natNT2-PGAL1-NDT80, spo11::KANMX, mps1-R170S::his5 |
| Y2187 | MAT∝, ura3::pAFS152[URA3 PCYC-GFP-lacI], trp1-Δ63, his3-Δ1, leu2, lys2::pMDE798[PDMC1-GFP-lacI], tyr1-2, met13-c, cyh2-1, SPC42-GFP-TRP1, natNT2-PGAL1-NDT80, PCLB2-3HA-NDC80 KanMX6, spo11::HIS3MX6 |
| X2298 | MAT∝, trp1-Δ63, his3-Δ1, leu2, met13-d, tyr1-1, lys2::pLL1[PCYC1-GFP-lacI LYS2], can1-R, ura3::pKB80 [PGPD1-GAL4(848)-ER-URA3::hphNT1], natNT2-PGAL1-NDT80, CEN1::pJN2[lacO256 LEU2], PCLB2-3HA-NDC80 KanMX6, spo11::HIS3MX6 |
| Y2188 | MATa, ura3::pAFS152[URA3 PCYC-GFP-lacI], trp1-Δ63, his3-Δ1, leu2, lys2::pMDE798[PDMC1-GFP-lacI], tyr1-2, met13-c, cyh2-1, SPC42-GFP-TRP1, natNT2-PGAL1-NDT80, PCLB2-3HA-NDC80 KanMX6, spo11::HIS3MX6 |
| X3217 | MATa, ura3-13, trp1-Δ63, his3-Δ1, leu2-?, met13-d, tyr1-1, lys2-1, can1-R, TUB1-OPL447[pHIS3p:mEos2-Tub1+3'UTR::TRP1] |
| Y2771 | MAT∝, leu2-?, lys2-2, met13-c, tyr1-2, ura3-1, trp1-Δ63, cyh2-1, his3-Δ1, TUB1-OPL447[pHIS3p:mEos2-Tub1+3'UTR::TRP1] |
| X3227 | MATa, ura3::pKB80 [PGPD1-GAL4(848)-ER-URA3::hphNT1], trp1-Δ63, his3-Δ1, leu2-?, met13-d, tyr1-1, lys2-1, can1-R, TUB1-OPL447[pHIS3p:mEos2-Tub1+3'UTR::TRP1], natNT2-PGAL1-NDT80 |
| Y2779 | MAT∝, leu2-?, lys2-2, met13-c, tyr1-2, ura3-1, trp1-Δ63, cyh2-1, his3-Δ1, TUB1-OPL447[pHIS3p:mEos2-Tub1+3'UTR::TRP1], natNT2-PGAL1-NDT80 |
| X3228 | MAT∝, ura3::pKB80 [PGPD1-GAL4(848)-ER-URA3::hphNT1], trp1-Δ63, his3-Δ1, leu2-?, met13-d, tyr1-1, lys2-1, can1-R, TUB1-OPL447[pHIS3p:mEos2-Tub1+3'UTR::TRP1], natNT2-PGAL1-NDT80 |
| Y2780 | MATa, leu2-?, lys2-2, met13-c, tyr1-2, ura3-1, trp1-Δ63, cyh2-1, his3-Δ1, TUB1-OPL447[pHIS3p:mEos2-Tub1+3'UTR::TRP1], natNT2-PGAL1-NDT80 |
| X3233 | MATa, ura3::pKB80 [PGPD1-GAL4(848)-ER-URA3::hphNT1], trp1-Δ63, his3-Δ1, leu2-?, met13-d, tyr1-1, lys2-1, can1-R, TUB1-OPL447[pHIS3p:mEos2-Tub1+3'UTR::TRP1], natNT2-PGAL1-NDT80, PCLB2-3HA-MPS1 KANMX6 |
| Y2783 | MAT∝, leu2-?, lys2-2, met13-c, tyr1-2, ura3-1, trp1-Δ63, cyh2-1, his3-Δ1, TUB1-OPL447[pHIS3p:mEos2-Tub1+3'UTR::TRP1], natNT2-PGAL1-NDT80, mps1Δ::KANMX, TRP1::10Xmyc-mps1-as1 |
| X3234 | MAT∝, ura3::pKB80 [PGPD1-GAL4(848)-ER-URA3::hphNT1], trp1-Δ63, his3-Δ1, leu2-?, met13-d, tyr1-1, lys2-1, can1-R, TUB1-OPL447[pHIS3p:mEos2-Tub1+3'UTR::TRP1], natNT2-PGAL1-NDT80, PCLB2-3HA-MPS1 KANMX6 |
| Y2784 | MATa, leu2-?, lys2-2, met13-c, tyr1-2, ura3-1, trp1-Δ63, cyh2-1, his3-Δ1, TUB1-OPL447[pHIS3p:mEos2-Tub1+3'UTR::TRP1], natNT2-PGAL1-NDT80, mps1Δ::KANMX, TRP1::10Xmyc-mps1-as1 |

| | |
|---|---|
| X3231 | *MATa, ura3::pKB80 [PGPD1-GAL4(848)-ER-URA3::hphNT1], trp1-Δ63, his3-Δ1, leu2-?, met13-d, tyr1-1, lys2-1, can1-R, TUB1-OPL447[pHIS3p:mEos2-Tub1+3'UTR::TRP1], natNT2-PGAL1-NDT80, spo11::KANMX* |
| Y2781 | *MAT∝, leu2-?, lys2-2, met13-c, tyr1-2, ura3-1, trp1-Δ63, cyh2-1, his3-Δ1, TUB1-OPL447[pHIS3p:mEos2-Tub1+3'UTR::TRP1], natNT2-PGAL1-NDT80, spo11::KANMX* |
| X3232 | *MAT∝, ura3::pKB80 [PGPD1-GAL4(848)-ER-URA3::hphNT1], trp1-Δ63, his3-Δ1, leu2-?, met13-d, tyr1-1, lys2-1, can1-R, TUB1-OPL447[pHIS3p:mEos2-Tub1+3'UTR::TRP1], natNT2-PGAL1-NDT80, spo11::KANMX* |
| Y2782 | *MATa, leu2-?, lys2-2, met13-c, tyr1-2, ura3-1, trp1-Δ63, cyh2-1, his3-Δ1, TUB1-OPL447[pHIS3p:mEos2-Tub1+3'UTR::TRP1], natNT2-PGAL1-NDT80, spo11::KANMX* |
| X3229 | *MATa, ura3::pKB80 [PGPD1-GAL4(848)-ER-URA3::hphNT1], trp1-Δ63, his3-Δ1, leu2-?, met13-d, tyr1-1, lys2-1, can1-R, TUB1-OPL447[pHIS3p:mEos2-Tub1+3'UTR::TRP1], natNT2-PGAL1-NDT80, spo11::KANMX, PCLB2-3HA-MPS1 KANMX6* |
| Y2787 | *MAT∝, leu2-?, lys2-2, met13-c, tyr1-2, ura3-1, trp1-Δ63, cyh2-1, his3-Δ1, TUB1-OPL447[pHIS3p:mEos2-Tub1+3'UTR::TRP1], natNT2-PGAL1-NDT80, spo11::KANMX, mps1-R170S::his5* |
| X3230 | *MATa, ura3::pKB80 [PGPD1-GAL4(848)-ER-URA3::hphNT1], trp1-Δ63, his3-Δ1, leu2-?, met13-d, tyr1-1, lys2-1, can1-R, TUB1-OPL447[pHIS3p:mEos2-Tub1+3'UTR::TRP1], natNT2-PGAL1-NDT80, spo11::KANMX, PCLB2-3HA-MPS1 KANMX6* |
| Y2805 | *MAT∝, leu2-?, lys2-2, met13-c, tyr1-2, ura3-1, trp1-Δ63, cyh2-1, his3-Δ1, TUB1-OPL447[pHIS3p:mEos2-Tub1+3'UTR::TRP1], natNT2-PGAL1-NDT80, mps1Δ::KANMX, TRP1::10Xmyc-mps1-as1, spo11::KANMX* |
| X3383 | *MATa, trp1-Δ63, his3-Δ1, leu2-?, met13-d, tyr1::[HIS5 pCUP1-AFB2], lys2-1, can1-R, STU2-AID*-9xMYC-HIS3MX6, ura3::pKB80 [PGPD1-GAL4(848)-ER-URA3::hphNT1], natNT2-PGAL1-NDT80, TUB1-OPL447[pHIS3p:mEos2-Tub1+3'UTR::TRP1]* |
| Y2933 | *MAT∝, leu2-?, lys2-2, met13-c, tyr1::[HIS5 pCUP1-AFB2], ura3-1, trp1-Δ63, cyh2-1, his3-Δ1, STU2-AID*-9xMYC-HIS3MX6, natNT2-PGAL1-NDT80, TUB1-OPL447[pHIS3p:mEos2-Tub1+3'UTR::TRP1]* |
| X3384 | *MAT∝, trp1-Δ63, his3-Δ1, leu2-?, met13-d, tyr1::[HIS5 pCUP1-AFB2], lys2-1, can1-R, STU2-AID*-9xMYC-HIS3MX6, ura3::pKB80 [PGPD1-GAL4(848)-ER-URA3::hphNT1], natNT2-PGAL1-NDT80, TUB1-OPL447[pHIS3p:mEos2-Tub1+3'UTR::TRP1]* |
| Y2934 | *MATa, leu2-?, lys2-2, met13-c, tyr1::[HIS5 pCUP1-AFB2], ura3-1, trp1-Δ63, cyh2-1, his3-Δ1, STU2-AID*-9xMYC-HIS3MX6, natNT2-PGAL1-NDT80, TUB1-OPL447[pHIS3p:mEos2-Tub1+3'UTR::TRP1]* |
| X1103 | *MATa, URA3::pKB80 [PGPD1-GAL4(848)-ER-URA3::hphNT1], trp1-63, his3-1, leu2, met13-d, tyr1-1, lys2::pLL1[PCYC1-GFP-lacI LYS2], can1 (R), ZIP1-TEV?, KanMX-PGAL1-NDT80, CEN1::pJN2[lacO256 LEU2]* |
| Y1092 | *MAT∝, ura3::pAFS152[URA3 PCYC-GFP-lacI], trp1-Δ63, his3-Δ1, leu2, lys2::pMDE798[PDMC1-GFP-lacI], tyr1-2, met13-c, cyh2-1, SPC42-[MDE1145: URA3 SPC42-DSRed], KanMX-PGAL1-NDT80, CEN1::pJN2[lacO256 LEU2]* |
| Y1124 | *MAT∝, ura3::pAFS152[URA3 PCYC-GFP-lacI], trp1-Δ63, his3-Δ1, leu2, lys2::pMDE798[PDMC1-GFP-lacI], tyr1-2, met13-c, cyh2-1, SPC42-[MDE1145: URA3 SPC42-DSRed], KanMX-PGAL1-NDT80, CEN1::pJN2[lacO256 LEU2], mps1Δ::KANMX, TRP1::10Xmyc-mps1-as1* |
